# Virophages and retrotransposons colonize the genomes of a heterotrophic flagellate

**DOI:** 10.1101/2020.11.30.404863

**Authors:** Thomas Hackl, Sarah Duponchel, Karina Barenhoff, Alexa Weinmann, Matthias G. Fischer

**Author notes:** Correspondence to Matthias G. Fischer, **Email:**.

## Abstract

Virophages can parasitize giant DNA viruses and may provide adaptive anti-giant-virus defense in unicellular eukaryotes. Under laboratory conditions, the virophage mavirus integrates into the nuclear genome of the marine flagellate *Cafeteria burkhardae* and reactivates upon superinfection with the giant virus CroV. In natural systems, however, the prevalence and diversity of host-virophage associations has not been systematically explored. Here, we report dozens of integrated virophages in four globally sampled *C. burkhardae* strains that constitute up to 2% of their host genomes. These <u>e</u>ndogenous <u>ma</u>virus-<u>l</u>ike <u>e</u>lements (EMALEs) separated into eight types based on GC-content, nucleotide similarity, and coding potential and carried diverse promoter motifs implicating interactions with different giant viruses. Between host strains, some EMALE insertion loci were conserved indicating ancient integration events, whereas the majority of insertion sites were unique to a given host strain suggesting that EMALEs are active and mobile. Furthermore, we uncovered a unique association between EMALEs and a group of tyrosine recombinase retrotransposons, revealing yet another layer of parasitism in this nested microbial system. Our findings show that virophages are widespread and dynamic in wild *Cafeteria* populations, supporting their potential role in antiviral defense in protists.

## Introduction

Many eukaryotic genomes harbor endogenous viral elements (EVEs) (1). For retroviruses, integration as a provirus is an essential part of their replication cycles, but other viruses also occasionally endogenize, for instance with the help of cellular retroelements (2). Some green algal genomes even contain giant EVEs of several hundred kilobase pairs (kbp) in length (3), but unlike prophages in bacteria and archaea, most eukaryotic EVEs are thought to be ‘genomic fossils’ and incapable of virion formation and horizontal transmission. However, some viral genes may be co-opted for various host functions (4, 5). In recent years, the exploration of protist-infecting giant viruses has uncovered a novel class of associated smaller DNA viruses with diverse and unprecedented genome integration capabilities.

Viruses of the family *Lavidaviridae*, commonly known as virophages, depend for their replication on giant DNA viruses of the family *Mimiviridae* and can parasitize them during coinfection of a suitable protist host (6–8). A striking example is the virophage mavirus, which strongly inhibits virion synthesis of the lytic giant virus CroV during coinfection of the marine heterotrophic nanoflagellate *Cafeteria* sp. (Stramenopiles; Bicosoecida) (9, 10). Virophages possess 15-30 kbp long double-stranded (ds) DNA genomes of circular or linear topology that tend to have low GC-contents (27-51%) (11). A typical virophage genome encodes 20-30 proteins, including a major capsid protein (MCP), a minor capsid or penton protein (PEN), a DNA packaging ATPase, and a maturation cysteine protease (PRO) (7). In addition to this conserved morphogenesis module, virophages encode DNA replication and integration proteins that were likely acquired independently in different virophage lineages (12). Viruses in the genus *Mavirus* contain a *rve*-family integrase (rve-INT) that is also found in retrotransposons and retroviruses, with close homologs among the eukaryotic Maverick/Polinton elements (MPEs) (9). MPEs were initially described as DNA transposons (13, 14), but many of them carry the morphogenesis gene module and thus qualify as endogenous viruses (15).

Phylogenetic analysis suggests that mavirus-type virophages share a common ancestry with MPEs and the related polinton-like viruses (PLVs) (9, 12). We therefore tested the integration capacity of mavirus using the cultured protist *C. burkhardae* (formerly *C. roenbergensis* (16, 17)) and found that mavirus integrates efficiently into the nuclear host genome (10). The resulting mavirus provirophages are transcriptionally silent unless the host cell is infected with CroV, which leads to reactivation and virion formation of mavirus. Newly produced virophage particles then inhibit CroV replication and increase host population survival during subsequent rounds of coinfection (10). The mutualistic *Cafeteria*-mavirus symbiosis may thus act as an adaptive defense system against lytic giant viruses (10, 18). The integrated state of mavirus is pivotal to the proposed defense scheme as it represents the host’s indirect antigenic memory of CroV (19). We therefore investigated endogenous virophages to assess the prevalence and potential significance of virophage-mediated defense systems in natural protist populations.

Here we report that mavirus-like EVEs are common, diverse, and most likely active mobile genetic elements (MGEs) of *C. burkhardae*. Our results suggest an influential role of these viruses on the ecology and evolution of their bicosoecid hosts.

## Results

### Endogenous virophages are abundant in *Cafeteria* genomes

In preparation of screening for endogenous virophages, we generated high-quality *de novo* genome assemblies of four cultured *C. burkhardae* strains (20). These strains, designated BVI, Cflag, E4-10P (E4-10) and RCC970-E3 (RCC970), were isolated from the Caribbean Sea in 2012, the Northwest Atlantic in 1986, the Northeast Pacific in 1989, and the Southeast Pacific in 2004, respectively. We sequenced their genomes using both short-read (Illumina MiSeq) and long-read (Pacific Biosciences RSII) technologies in order to produce assemblies that would resolve 20-30 kb long repetitive elements within the host genomic context. Each *C. burkhardae* genome assembly comprised 34-36 megabase pairs with an average GC-content of 70% (20).

To identify endogenous virophages, we combined sequence similarity searches against known virophage genomes with genomic screening for GC-content anomalies. The two approaches yielded redundant results and virophage elements were clearly discernible from eukaryotic genome regions based on their low (30-50%) GC-content (**Fig. 1A**). Each element had at least one open reading frame (ORF) with a top blastp hit to a mavirus protein, with no elements bearing close resemblance to Sputnik or other virophages outside the genus *Mavirus*. In the four *Cafeteria* genomes combined, we found 138 <u>e</u>ndogenous <u>ma</u>virus-like <u>e</u>lements (EMALEs, **Figs. 1B,C**, S1, Table S1). Thirty-three of these elements were flanked by terminal inverted repeats (TIRs) and host DNA and can thus be considered full-length viral genomes.

**Figure 1.**
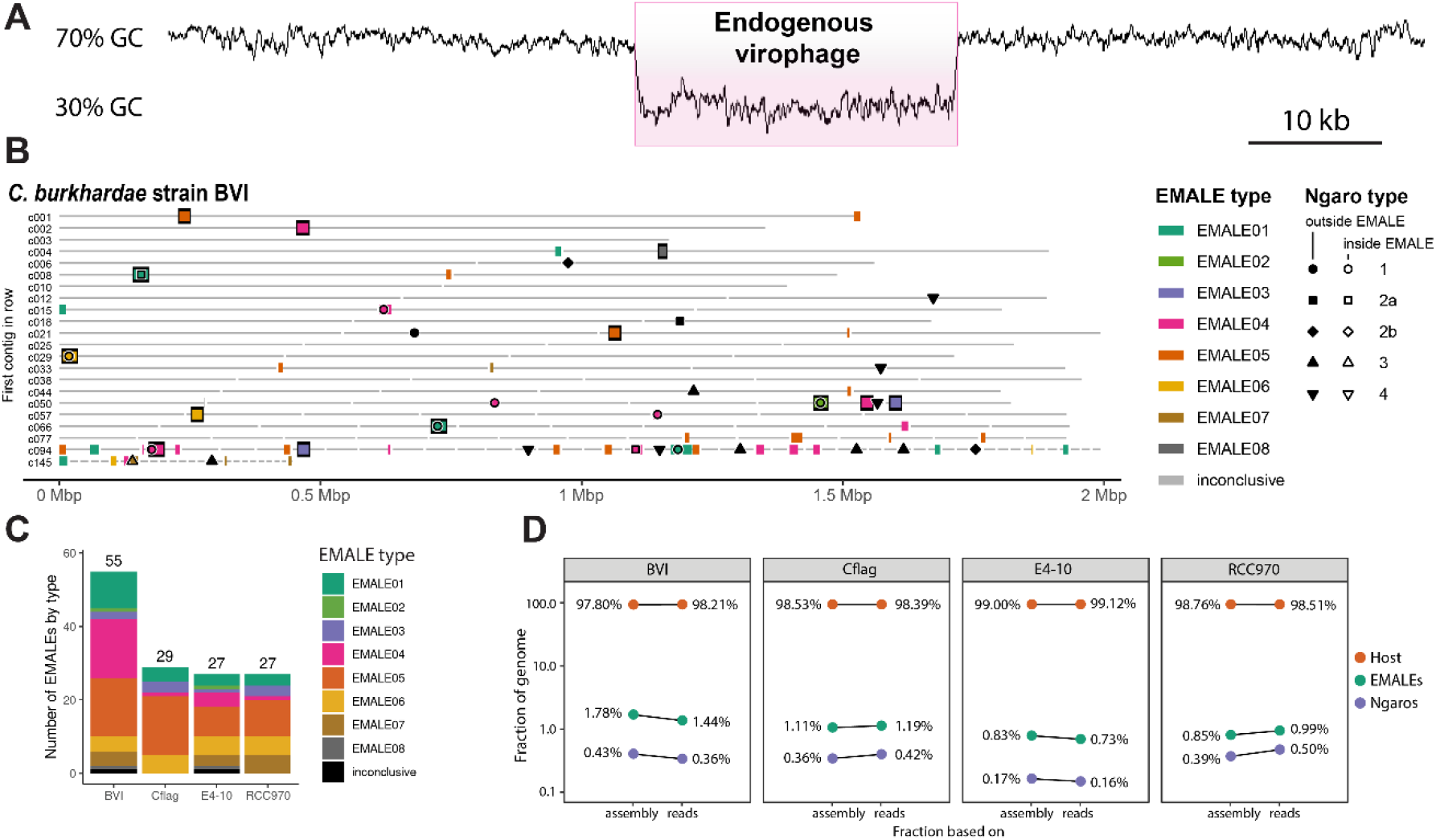
Endogenous virophages in *Cafeteria burkhardae*. (A) GC-content graph signature of a virophage element embedded in a high-GC host genome. Shown is a region of contig BVI_c002 featuring an integrated virophage (pink box) flanked by host sequences. (B) Location of partial or complete virophage genomes and Ngaro retrotransposons in the genome assemblies of *C. burkhardae* strain BVI (see Fig. S1 for all four strains). Horizontal lines represent contigs in order of decreasing length with numbers shown for the first contig of each line; colored boxes indicate endogenous mavirus-like elements (EMALEs). Fully assembled elements are framed in black. Ngaro retrotransposon positions are marked by black symbols; open symbols indicate Ngaros integrated inside a virophage element.

The remainder were partial virophage genomes that were located at contig ends or on short contigs. In these cases, the assembly algorithm probably terminated due to the presence of multiple identical or highly similar EMALEs within the same host genome – a well-known issue for repetitive sequences (21). With 55 elements, *C. burkhardae* strain BVI contained nearly twice as many EMALEs as any of the other strains, where we found 27-29 elements per genome (**Fig. 1C**, Table S1). Compared to the total assembly length, EMALEs composed an estimated 0.7% to 1.8% of each host assembly (**Fig. 1D**). Contributions calculated from assemblies deviated only slightly (0.01%-0.3%) from read-based calculations. Therefore, the assemblies seem to provide a good representation of the actual contribution of EMALEs to the overall host genomes.

### Endogenous mavirus-like elements (EMALEs) are genetically diverse

From here on, we focus our analysis on the 33 complete virophages genomes, which were 5.5 kb to 21.5 kb long with a median length of 19.8 kb, and TIRs that varied in length from 0.2 kb to 2.3 kb with a median of 0.9 kb (Table S1, Fig. S2). Their GC-contents ranged from 29.7% to 52.7%, excluding retrotransposon insertions where present. To classify EMALEs we used an all-versus-all DNA dot plot approach (**Fig. 2**). It revealed two main blocks: The first block contained EMALEs with GC-contents of 29.7% to 38.5% (median 35.3%), whereas EMALEs in the second block had GC-contents ranging from 47.2% to 52.7% (median 49.3%). The *C. burkhardae* EMALEs can thus be roughly separated into low-GC and mid-GC groups. Based on the similarity patterns within each block, we further distinguish eight EMALE types, with low-GC EMALEs comprising types 1-4 and mid-GC EMALEs comprising types 5-8 (**Fig. 2**). Representative genome diagrams for each EMALE type are shown in **Fig. 3**, for a schematic of all 33 complete EMALEs see Fig. S2. According to this classification scheme, the reference mavirus strain Spezl falls within type 4 of the low-GC EMALEs (**Figs. 2, 3**, S2). Partial EMALEs were classified based on their sequence similarity to full-length type species (Fig. S3).

**Figure 2.**
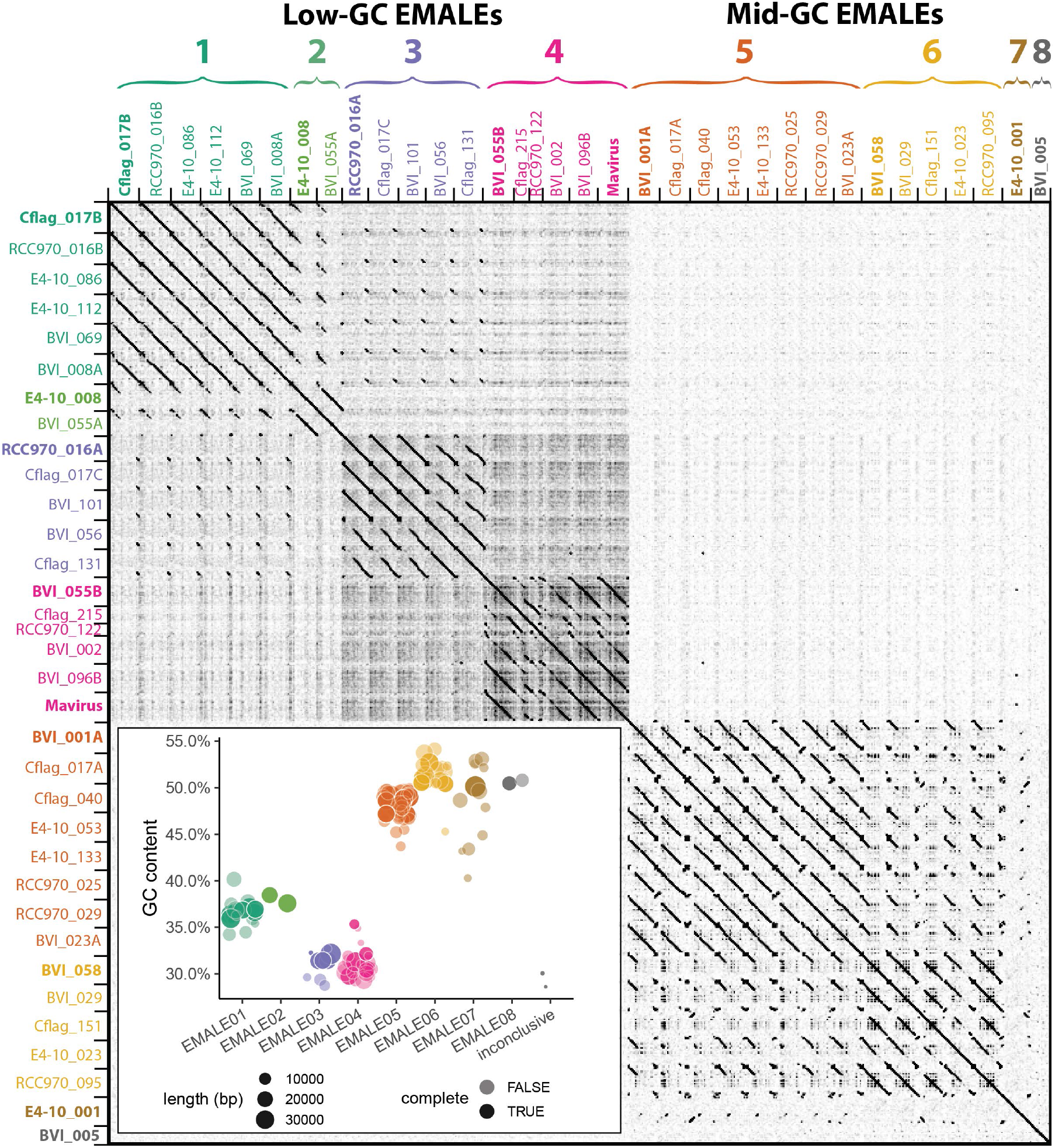
Classification of endogenous virophages based on DNA dot plot analysis. The self-versus-self DNA dot plot of concatenated sequences of 33 complete EMALE genomes and mavirus reveals two main block patterns, corresponding to EMALEs with low (29%-38%) GC-content and medium (47%-53%) GC-content. Smaller block patterns define EMALE types 1-8. EMALE identifiers indicate the host strain and contig number where the respective element is found. Multiple EMALEs on a single contig are distinguished by terminal letters. Elements printed in bold represent the type species shown in Fig. 3. Inset: GC-content distribution of complete and partial EMALEs. Some partial EMALEs were too short for type assignment and are thus inconclusive. Retrotransposon insertions, where present, were removed prior to analysis.

**Figure 3.**
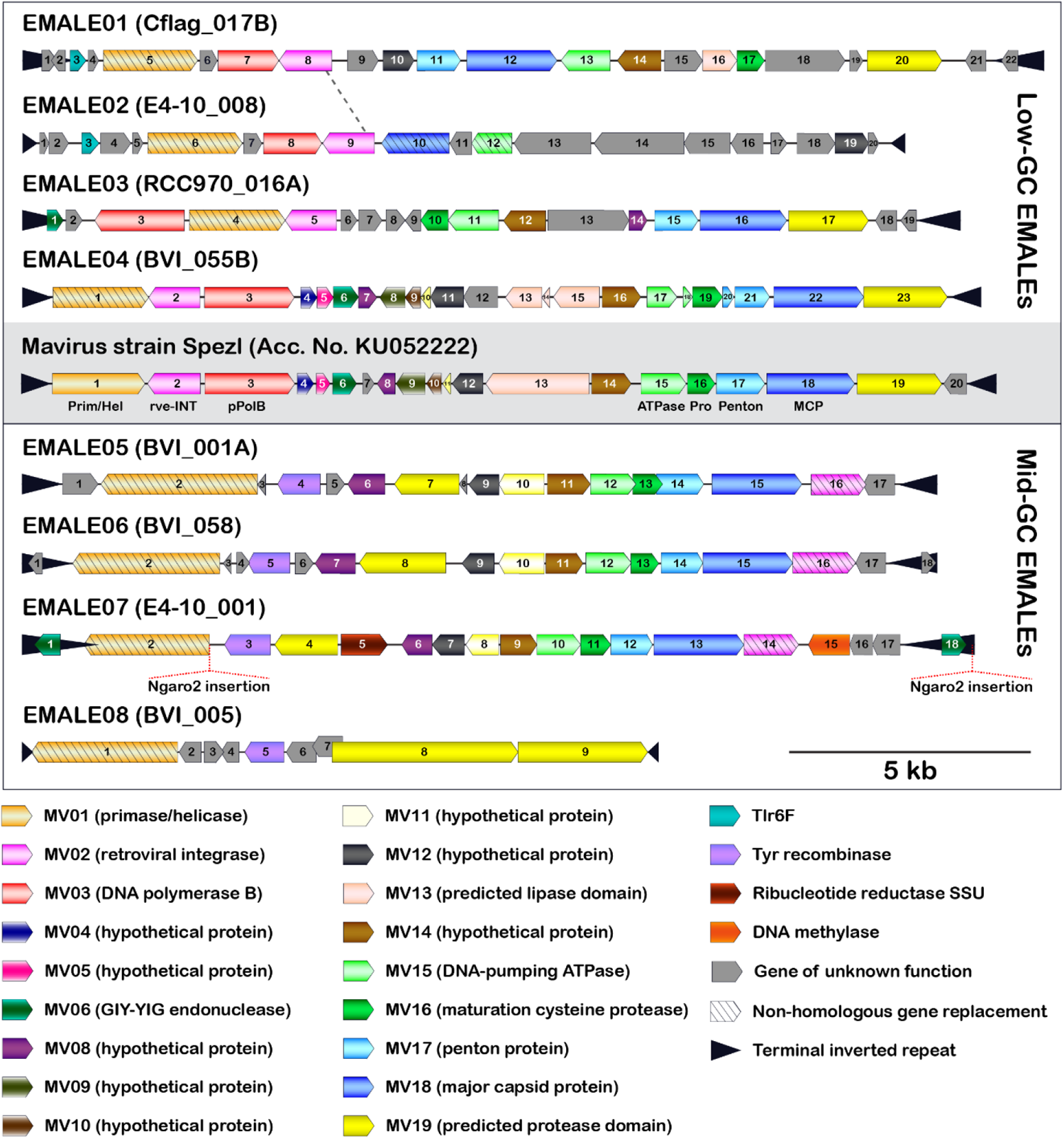
Genome organization of eight EMALE types found in *C. burkhardae*. Shown are schematic genome diagrams of the EMALE type species 1-8; for all 33 complete EMALEs see Fig. S2. The reference mavirus genome with genes MV01-MV20 is included for comparison. Homologous genes are colored identically; genes sharing functional predictions but lacking sequence similarity to the mavirus homolog are hatched. Open reading frames are numbered individually for each element. Ngaro retrotransposon insertion sites are indicated where present. The dotted line between EMALE01 and EMALE02 separates a homologous region (left) from unrelated DNA sequences (right) and thus indicates the location of a probable recombination event.

The codon and amino acid composition of EMALE genes clearly correlated with the overall GC-content of the EMALE genomes (Fig. S4). For each encoded amino acid, we observed a strong shift towards synonymous codons reflecting the overall GC trend, and across amino acids, we observed a shift from those encoded by high-GC codons to those encoded by low-GC codons in low-GC EMALEs and vice versa. This uniform trend across all amino acids likely indicates that selection and evolutionary processes driving GC-content variation in these viruses act on the nucleotide level, rather than on the encoded proteins.

With few exceptions, EMALEs are predicted to encode 17-21 proteins each. None of the encoding genes was found to contain introns. The virion morphogenesis module in EMALE types 1 and 3-7 consists of the canonical virophage core genes corresponding to MCP, PEN, ATPase, and PRO proteins. Type 2 EMALEs likely encode a different set of capsid genes as discussed below, and the truncated EMALE type 8 lacks recognizable morphogenesis genes. Another highly conserved gene in EMALE types 1 and 3-7 is *MV14*, which is always found immediately upstream of the *ATPase* (**Figs. 3**, S2) and codes for a protein of unknown function that is part of the mavirus virion (22). *MV14* is present in various metagenomic virophage sequences (23) and, based on synteny and protein localization, likely encodes an important virion component in members of the genus *Mavirus*.

The replication/integration module consists of the *rve-INT* gene and at least one additional ORF coding for a primase/helicase and a DNA polymerase. Low-GC EMALEs encode a mavirus-related primase/helicase and protein-primed family B DNA polymerase (pPolB) (**Figs. 3**, S2). Mid-GC EMALEs, on the other hand, lack the *pPolB* gene and feature a longer primase/helicase ORF that may include a DNA polymerase domain similar to the helicase-polymerase fusion genes described in PLVs (24).

Other mavirus genes frequently found in EMALEs include *MV19* (encoding a putative protease domain), and two genes of unknown function, *MV08* and *MV12*. Interestingly, all mid-GC EMALEs encode a predicted tyrosine recombinase (YR) in addition to the *rve-INT* and thus possess two predicted enzymes for genome integration. YRs have been found in other virophages and likely catalyze integration into giant virus genomes (25, 26). Notable genes unique to one EMALE type include a putative DNA methylase and a ribonucleotide reductase small subunit gene found in EMALE07. The Tlr6F protein encoded by EMALE types 1+2 is present in diverse MGEs, including other virophages, PLVs, and large DNA viruses of the phylum *Nucleocytoviricota* (19, 27).

In general, genes were syntenic between EMALEs of the same type, whereas gene order was poorly conserved among EMALEs of different types, with the following exceptions: *MCP* was always preceded by *PEN*, and *ATPase* was always preceded by *MV14*, whereas the *MV14-ATPase-PRO-PEN-MCP* morphogenesis gene order as seen in mavirus was present only in EMALE types 4-7. EMALE02 represents an interesting case, as it shares 6-7 kb of its 5’ part (we chose the *primase/helicase* genes to mark the 5’ end of all EMALEs) with EMALE01, while the remaining 11 kb are not closely related to other EMALEs or virophages (Fig. S5). Genes encoded in the latter region are mostly ORFans, with the exception of a *MV12*-like gene and divergent *MCP* and *ATPase* genes with remote similarity to PLVs (28) and adintoviruses (29). EMALE02 may thus be the result of a recombination event that exchanged the canonical virophage morphogenesis module of EMALE01 with capsid genes of a PLV (**Fig. 3**, dashed line). Overall, these observations support the notion that recombination and non-homologous gene replacement are important factors in virophage genome evolution (12).

### Core gene conservation and non-homologous gene replacement in EMALEs

To validate our classification scheme for EMALEs and to place them in a phylogenetic context to other virophages, we used maximum likelihood reconstruction on the core proteins MCP, PEN, ATPase, and PRO, as well as on rve-INT (**Fig. 4**). In the resulting phylogenetic trees, EMALE core proteins formed monophyletic clades with mavirus and related sequences from environmental samples, thus significantly expanding the known diversity of the genus *Mavirus*. The environmental sequences that clustered with EMALE core proteins include a single amplified genome (SAG) from an uncultured chrysophyte (30), the metagenomic Ace Lake Mavirus (ALM) (31), and four additional metagenomes that were identified in a global survey of virophage sequences (23). The chrysophyte SAG is nearly identical to mavirus strain Spezl and indicates that the host range of mavirus extends beyond bicosoecids. The metagenomic sequences either clustered with one of the EMALE types, or branched separately from them, which suggests the existence of additional sub-groups (e.g. M590M2_1006461).

**Figure 4.**
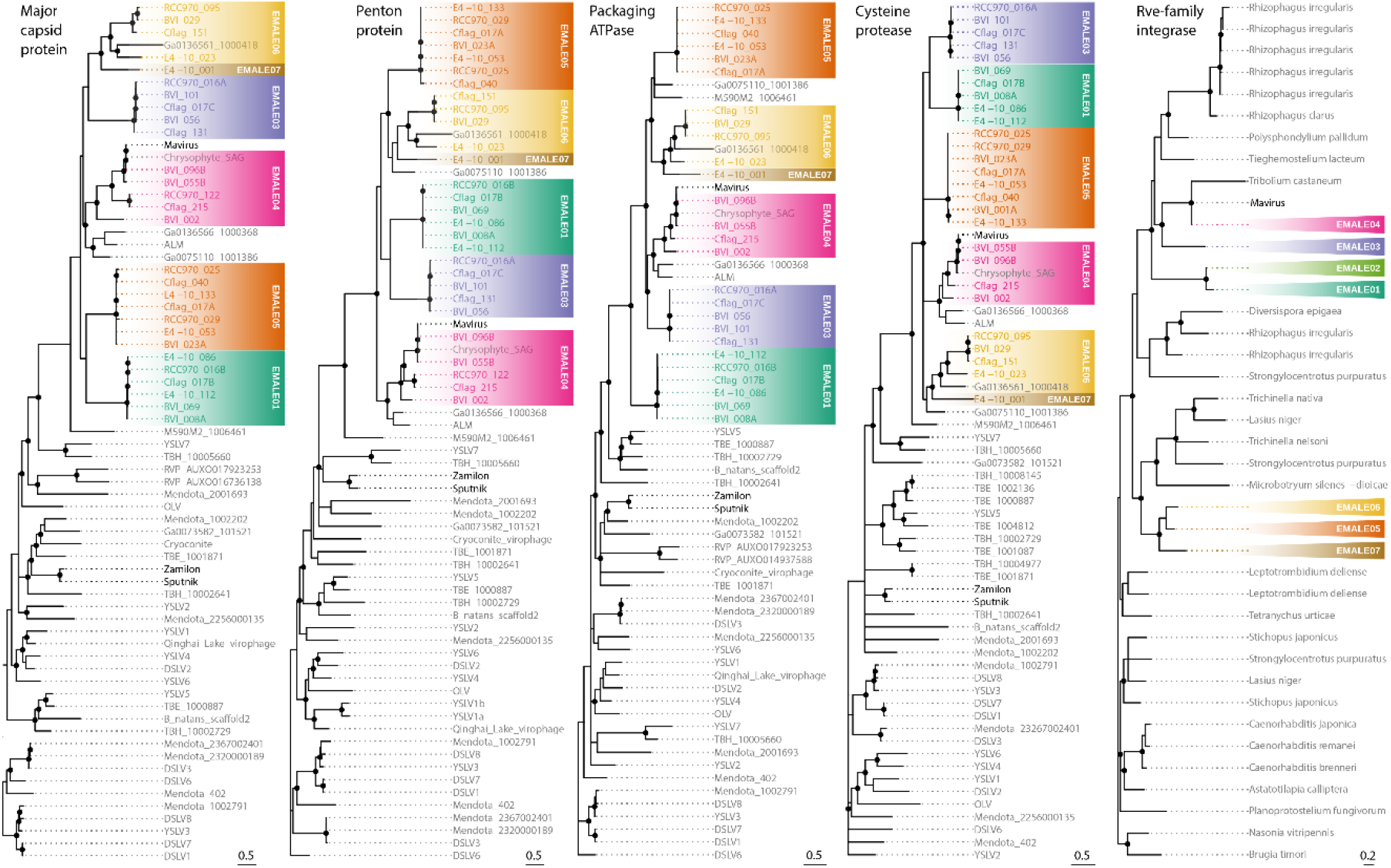
Phylogenetic reconstruction of conserved EMALE proteins. Unrooted maximum likelihood trees were constructed from multiple sequence alignments of the four virophage core proteins MCP, PEN, ATPase, and PRO, as well as of the retroviral integrase. Nodes with bootstrap values of 80% or higher are marked with dots. EMALEs are color-coded by type; cultured virophages are printed in bold. ALM, Ace Lake Mavirus; DSLV, Dishui Lake virophage; OLV, Organic Lake virophage; RVP, rumen virophage; TBE/TBH, Trout Bog Lake epi-/hypolimnion; YSLV, Yellowstone Lake virophage. Metagenomic sequences starting with Ga and M590 are derived from (23).

Within the *Mavirus* clade, EMALEs of a given type were monophyletic for each of the four core proteins, which corroborates their dot plot-based classification. It is worth noting that although EMALEs of type 5 and 6 are largely syntenic (**Figs. 3**, S2), they were clearly distinguishable in their phylogenetic signatures (**Fig. 4**). A comparison of clade topologies revealed that even within the conserved morphogenesis module, individual proteins differed with regard to their neighboring clades, and low-GC and mid-GC EMALEs did not cluster separately from each other. These observations could suggest that the morphogenesis modules of different EMALE types diversified simultaneously and that adaptation of GC-content may occur rather quickly.

In contrast, phylogenetic analysis of rve-INT proteins revealed separate clades for low-GC and mid-GC EMALEs (**Fig. 4**). Each of these clades was affiliated with different cellular homologs that included MPEs and retroelements. Notably, the *rve-INT* genes of low-GC EMALEs were located near the 5’ end of the virophage genome, whereas in mid-GC EMALEs, they were located near the 3’ end (**Figs. 3**, S2). These observations suggest that EMALEs encode two different rve-INT versions, one specific for low-GC EMALEs that co-occurs with the pPolB and a shorter primase/helicase ORF, and one specific for mid-GC EMALEs that co-occurs with a longer primase/helicase ORF. The two integrase versions may have been acquired independently, or one version could have replaced the other during EMALE evolution.

Such non-homologous gene replacement appears to have taken place among the primase/helicase genes, too, as previously noted for virophages in general (12). EMALEs encode several different versions of primase/helicase genes with a degree of amino acid divergence that precluded their inclusion in a single multiple sequence alignment. The YR proteins encoded by EMALE types 5-8 formed a monophyletic clade and were part of a larger group of recombinases that included virophages from freshwater metagenomes, as well as microalgae and algal nucleocytoviruses (Fig. S6).

### *Cafeteria* strains differ in their virophage composition

The four *C. burkhardae* strains displayed distinct EMALE signatures: strain BVI had the highest number of virophage elements with 13 complete and 42 partial EMALEs, whereas the other three strains had 6-7 complete and 20-22 partial EMALEs each (**Fig. 1C**, Table S1). EMALE types 1,3,4,5, and 6 were present in every host strain, EMALE07 was found in all strains except Cflag, and EMALE types 2 and 8 were detected in strains BVI and E4-10 only. We found no evidence for sequence-specific genome integration of EMALEs after inspecting the host DNA sequences that flanked EMALE integration sites, which confirms previous reports of mavirus integration (10). EMALEs were flanked by target site duplications (TSDs) that were predominantly 3-5 bp in length, although some were as short as 1 bp or as long as 9 bp (Table S2). By comparison, mavirus and MPEs generate 5-6 bp long TSDs upon integration (10, 13, 14).

To assess whether homologous EMALEs were found in identical loci in closely related host genomes, we conducted sequence similarity searches with the flanking regions of each of the 33 fully resolved EMALEs. Whenever these searches returned a homologous full or partial EMALE with at least one matching host flank, we considered the EMALE locus to be conserved in these host strains. We found varying degrees of conservation, with examples shown in Fig. S7. In 11 cases, an EMALE insertion was conserved in at least two host strains (Table S2): three EMALE loci were shared by all four strains, four were shared by three strains, and another four were shared by two strains. Based on conserved EMALE loci, strains Cflag and RCC970 were most closely related with nine shared EMALE integrations, which is in line with phylogenetic and average nucleotide identity (ANI) analyses of these strains (20). The four *C. burkhardae* genomes have ANIs of >99% and thus appear to differ mostly based on their content of EMALEs and other MGEs.

The most parsimonious scenario for the origin of EMALEs that are located in identical loci in different host strains is that they derived from a single integration event. For instance, EMALE03 BVI_101 is orthologous to Cflag_017C and RCC970_016A (Fig. S7C), which suggests that this element initially colonized the common ancestor of *C. burkhardae* strains BVI, Cflag, and RCC970. Further cases of redundant EMALEs are Cflag_017B & RCC970_016B (EMALE01) and BVI_029 & RCC970_095 (EMALE06). These elements may thus derive from relatively ancient integration events, whereas 18 of the 33 complete EMALEs represent integrations that were unique to a single host strain (Table S2). Strain BVI contained 10 of these 18 unique integrations, more than twice as many as any other strain.

The genomic landscape around EMALE integration sites ranged from repeat-free flanking regions to complex host repeats (Fig. S8). Of the 29 different integration sites represented by the 33 fully resolved EMALEs, 18 were located near repetitive host DNA (within 10 kb from the insertion site). These repeats, in addition to EMALE TIRs, multiple copies of the same EMALE type, and the putative heterozygosity of EMALE insertions, occasionally caused assembly problems, as illustrated in Fig. S7.

Next, we analyzed whether EMALE insertions interrupted coding sequences of the host. Fifteen integration sites were located within a predicted host gene (13 in exons, 2 in introns), four were found in predicted 3’ untranslated regions, and three were located in intergenic regions (Table S2). These data show that EMALE insertions may disrupt eukaryotic genes with potential negative consequences for the host. The apparent preference for integration in coding regions could be assembly related, driven by increased accessibility of euchromatin, or linked to host factors that could direct the rve-INT via its CHROMO domain (32).

### EMALEs are predicted to be functional and mobile

Based on genomic features such as coding potential, ORF integrity, and host distribution, most EMALEs appear to be active MGEs. With the exception of EMALE08 and EMALE02, all endogenous *Cafeteria* virophages encode the canonical morphogenesis gene module consisting of *MCP, PEN, ATPase, PRO*, as well as *MV14*. EMALE02 likely encodes more distantly related capsid genes. Therefore, all EMALE types except EMALE08 should be autonomous for virion formation. In addition, all EMALEs contain at least one predicted enzyme for genome integration, an rve-INT in EMALE types 1-7 and a YR in EMALE types 5-8. EMALEs thus encode the enzymatic repertoire for colonizing new host genomes. Finally, the high variability of EMALE integration loci among otherwise closely related host strains strongly argues for ongoing colonization of natural *Cafeteria* populations by virophages.

The genomic similarity to mavirus implies that EMALEs may also depend on a giant virus for activation and horizontal transmission. Shared regulatory sequences in virophages and their respective giant viruses suggest that the molecular basis of virophage activation lies in the recognition of virophage gene promoters by giant virus encoded transcription factors (9, 33, 34). We therefore analyzed the 100 nt upstream regions of EMALE ORFs for conserved sequence motifs using MEME (35). For all type 4 EMALEs, which include mavirus, we recovered the previously described mavirus promotor motif ‘TCTA’, flanked by AT-rich regions. This motif corresponds to the conserved late gene promoter in CroV (9, 36), thus possibly indicating that all type 4 EMALEs could be reactivated by CroV or close relatives. EMALEs of other types lacked the ‘TCTA’ motif, but contained putative promoter sequences that may be compatible with different giant viruses (Fig. S9).

MGEs are prone to various decay processes including pseudogenization, recombination, and truncation. Among the 33 fully resolved EMALEs are three truncated elements: Cflag_215 and RCC970_122 (both EMALE04), and BVI_005 (EMALE08) (Fig. S2). Interestingly, even these shorter elements are flanked by TIRs, which must have regenerated after the truncation event. Whereas most EMALE ORFs appeared to be intact, as judged by comparison with homologous genes on syntenic elements, several EMALEs contained fragmented ORFs (e.g. *ATPase* and *PEN* genes in EMALE04 BVI_055B, **Fig. 3**). To test whether premature stop codons may be the result of assembly artifacts, we amplified selected EMALEs by PCR and analyzed the products using Sanger sequencing. When we compared the Sanger assemblies with the Illumina/PacBio assemblies, we noticed that the latter contained several substitutions and indels. For example, the MCP gene of EMALE01 RCC970_016B was split into three ORFs in the Illumina/PacBio assembly, whereas a single ORF was present in the corresponding Sanger assembly (Fig. S10). None of the Sanger assemblies confirmed fragmentation of conserved virophage genes, emphasizing the importance of independent sequence validation. However, since it was not possible to re-sequence all potential pseudogene locations, we cannot exclude that some EMALE genes may be fragmented. Overall, EMALE ORFs appeared to be intact and are thus likely to encode functional proteins.

### Tyrosine recombinase retrotransposons integrate into EMALEs

Strikingly, about one in four virophage elements was interrupted by a GC-rich sequence with a typical length of ∼6 kb (**Fig. 5A**, Table S1). We identified these insertions as retrotransposons of the Ngaro superfamily, within the DIRS order of retrotransposons (37). Ngaro retrotransposons feature split direct repeats with A_1_–[ORFs]–B_1_A_2_B_2_ structure (**Fig. 5D**), encode a YR instead of the rve-INT that is typical for retrotransposons, and are found in various eukaryotes from protists to vertebrates (38). In the four *C. burkhardae* strains, we annotated 80 Ngaro elements, with 10 to 25 copies per strain (Table S3). In addition, we found isolated AB repeats scattered throughout the genome, which could have arisen from recombination of 5’ and 3’ repeats and are reminiscent of solo long terminal repeats of endogenous retroviruses (39). Dot plot analysis of concatenated *Cafeteria* Ngaro sequences revealed four distinct types that showed no similarity at the nucleotide level, but appeared to share the same coding potential (**Fig. 5B,D**). Based on synteny to previously described Ngaros, ORF1 may encode a Gag-like protein; ORF2 encodes a predicted reverse transcriptase and ribonuclease H domains; and ORF3 encodes a predicted YR with the conserved His-X-X-Arg motif and catalytic Tyr residue. Ngaro YRs are related to putative transposons of bacteria and eukaryotes (Fig. S11), but bear no sequence similarity to the EMALE-encoded YRs.

**Figure 5.**
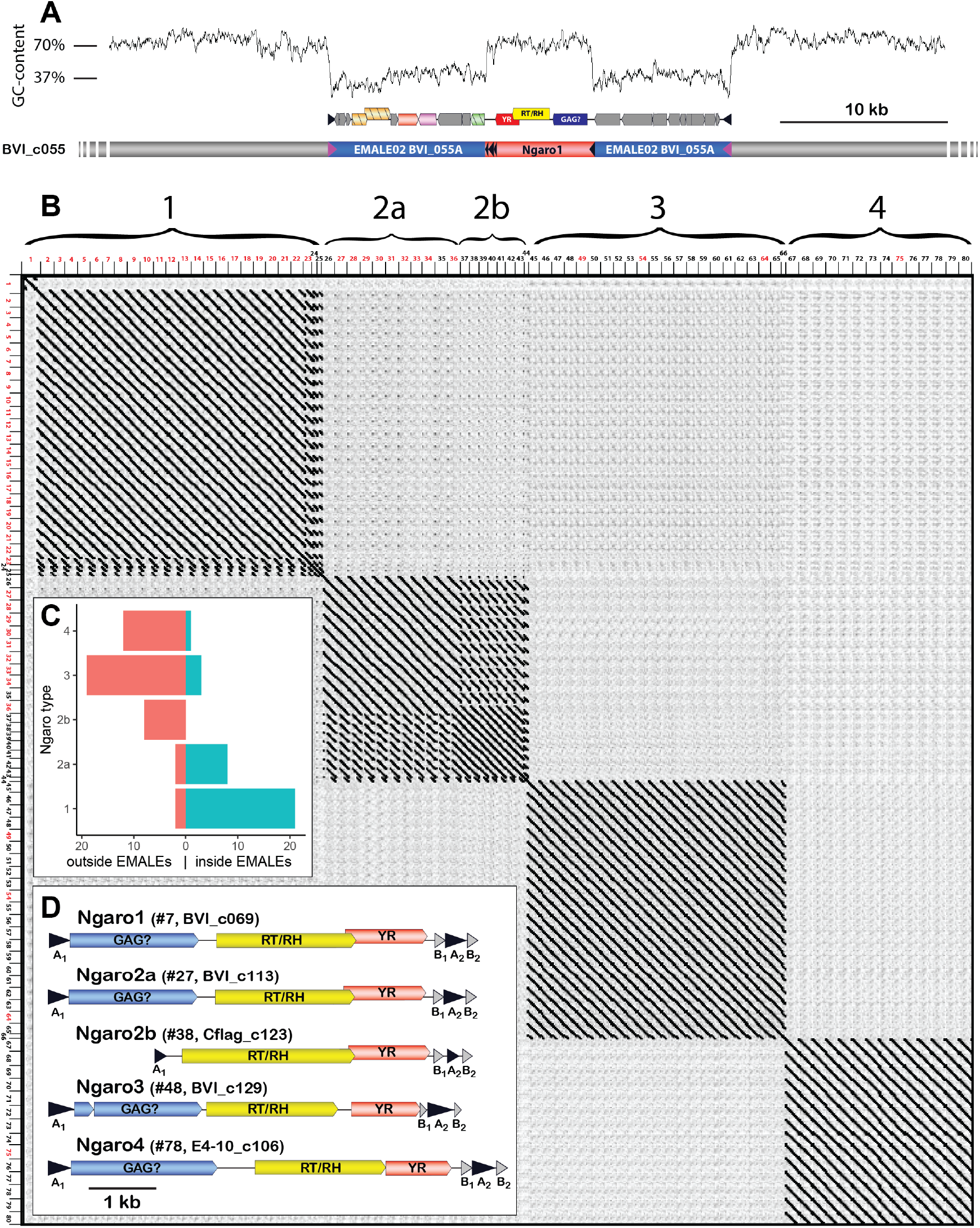
Ngaro retrotransposons in *C. burkhardae*. (A) Genomic profile of an EMALE-integrated Ngaro element showing a GC-content graph (top), ORF organization of EMALE and Ngaro (middle) and a schematic overview of the three genomic entities (bottom; host: grey, EMALE: blue, Ngaro: red). (B) Self-versus-self DNA dot plot of 80 concatenated Ngaro sequences. Block patterns define Ngaro types 1-4. Ngaros are numbered according to Table S3, with red numbers indicating retrotransposons inserted in EMALEs. (C) Distribution of Ngaro integration loci in EMALE and host DNA. Ngaro types 1 and 2a show a clear preference for EMALE loci, in contrast to Ngaro types 2b, 3, and 4 that are mostly found in host loci. (D) Coding potential of *C. burkhardae* Ngaro retrotransposons, shown for one example per type with their host strain and contig numbers listed. Triangles indicate direct repeats. GAG, group specific antigen; RT, reverse transcriptase; RH, ribonuclease H; YR, tyrosine recombinase.

Interestingly, the four Ngaro types in *C. burkhardae* differed with regard to their integration site preference. Whereas 91% of type 1 Ngaros were inserted in an EMALE, this was the case for only 14% and 8% of type 3 and type 4 Ngaros, respectively. At first glance, type 2 Ngaros were distributed evenly among viral and eukaryotic DNA (47% inside EMALEs, 53% outside EMALEs). However, several type 2 Ngaros were truncated at their 5’ end, featuring ∼2 kb long deletions that covered ORF1. All of these deletion variants, which we designate as type 2b, were located in host DNA, in contrast to 82% of the full-length type 2a Ngaros that were inserted in EMALEs (**Fig. 5C,D**). Similarly, most of the type 1 Ngaros that were inserted into eukaryotic chromatin also lacked ORF1. Hence, it appears that ORF1 determines the integration site specificity of Ngaro retrotransposons. Out of 20 Ngaro insertion sites with analyzable EMALE flanking regions, 9 were located in intergenic regions, 7 in EMALE genes, and 4 in TIRs (Table S3). Considering that intergenic regions comprise only 5-10% of an EMALE genome, we notice a significant bias (p<1e^-5^, Fisher’s exact test) towards Ngaro integration in intergenic EMALE DNA, which may be caused either by purifying selection of deleterious Ngaro insertions, or by a higher preference for Ngaro integration into EMALE intergenic regions due to their lower GC-content.

A possible consequence of retrotransposon insertion is the loss of biological activity and subsequent pseudogenization. However, we found that Ngaro-containing EMALEs did not contain more fragmented genes than Ngaro-free EMALEs (Fig. S12). Whereas the biological properties of Ngaro retrotransposons and their influence on host-virus-virophage dynamics remain to be explored, the EMALE-Ngaro interactions appear to be convoluted. For instance, an EMALE03 genome in strain Cflag is interrupted by two adjacent Ngaro1 insertions, while the EMALE itself is located inside an Ngaro4 element (Fig. S13).

## Discussion

Virophages represent a recently discovered family of eukaryotic dsDNA viruses that possess interesting genome integration properties and have potentially far-reaching eco-evolutionary consequences. Our genomic survey of the marine bicosoecid *Cafeteria burkhardae* revealed an unexpected abundance and diversity of endogenous virophages, with dozens of elements in a single host genome. Based on DNA dot plots and phylogenetic analysis, we distinguish eight different types of mavirus-related endogenous virophages. Similar to mavirus, these EMALEs could potentially reactivate and replicate in the presence of a compatible giant virus. Mavirus is proposed to act as an adaptive defense system against CroV in *Cafeteria* populations (10, 18), and our findings suggest that different types of EMALEs may respond to different giant viruses infecting *Cafeteria*. The assortment of endogenous virophages in a given host genome may thus reflect the giant virus infection history of that population (40). Some EMALEs are present in orthologous genomic loci in two or more host strains and likely date back to the common ancestor of these strains. However, at least half of the EMALE insertions are specific to a given host strain and may thus have been acquired relatively recently. Combined with the overall integrity of EMALEs and the conservation of integrase and capsid genes, these findings suggest that endogenous virophages in *C. burkhardae* are active MGEs.

Horizontal transmission of virophages in natural environments is likely limited by their requirement for two concurrent biological entities, namely a susceptible host cell infected with a permissive giant virus. Similar to temperate bacteriophages, persistence in the proviral state may thus be an essential survival strategy for virophages, as underscored by the abundance of EMALEs in *C. burkhardae*. Endogenous virophages may also be common in eukaryotes outside the order Bicosoecales, although a search for provirophages in 1,153 eukaryotic genomes found only one clear case in the chlorarachniophyte alga *Bigelowiella natans* (40). We propose that the discovery of host-integrated virophages is hampered not only by sampling bias, but also by technical limitations. For instance, AT-rich mavirus DNA was severely underrepresented when we sequenced the genome of *C. burkhardae* strain E4-10M1 with the standard Illumina MiSeq protocol (10). Additional problems arise during binning and assembly procedures. Although our sequencing and assembly strategy was specifically tailored to endogenous virophages and resolved 33 EMALEs in their host genomic context, dozens of EMALEs were only partially assembled, some may contain assembly errors (Fig. S7), and others may have been missed altogether. Advances in long-read sequencing technologies and assembly algorithms will likely alleviate such problems.

Surprisingly, we found that EMALEs were frequently interrupted by Ngaro retrotransposons, which revealed an additional level of nested parasitism in this microbial system. *Cafeteria* genomes contain four distinct Ngaro types with different affinities for EMALEs. Deletion of ORF1 in type 1 and type 2 Ngaros coincides with a decreased occurrence of these retrotransposons in EMALEs (**Fig. 5**). Syntenic ORFs in other Ngaros are predicted to encode a Gag-like structural protein (41), and Gag proteins of several retroviruses have been linked to integration site specificity (42, 43). The putative Gag proteins of *Cafeteria* Ngaros may thus influence whether retrotransposon insertion occurs in an EMALE or in eukaryotic chromatin. So far, retrotransposons have not been described for giant DNA viruses or virophages; however, pandoravirus genomes contain DNA transposons (44), and a class of 7 kb-long DNA MGEs called transpovirons interacts with the particles and genomes of *Acanthamoeba*-infecting mimiviruses and their virophages, apparently without affecting viral replication (25, 45). It remains to be studied whether Ngaro retrotransposons use reactivated virophages or giant viruses as vehicles for horizontal transmission, and what effect retrotransposon insertion in EMALEs has on the fitness of the virophage, the host cell, and their associated giant viruses.

In conclusion, we show that endogenous virophage genomes are abundant and diverse in the marine heterotrophic protist *Cafeteria burkhardae*. These mavirus-like EVEs appear to be active and dynamic MGEs with significant potential to shape the genome evolution of their hosts. We present evidence for recombination and gene exchange within EMALEs, and a previously unknown affiliation between virophages and YR retrotransposons. Our findings imply an important role for EMALEs in the ecology and evolution of bicosoecids and are in line with the hypothesis that endogenous virophages provide adaptive defense against giant viruses.

## Materials and Methods

### *Cafeteria burkhardae* cultures & genome sequencing

*C. burkhardae* strains BVI, Cflag, E4-10P, and RCC970-E3 were cultured, and their genomes sequenced & assembled as described previously (20).

### Nucleotide contributions of EMALEs and Ngaros to *Cafeteria* genomes

To quantify how much of each host genome is comprised of EMALEs and Ngaros, we applied two complementary strategies: i) We compared the number of nucleotides annotated as EMALEs and Ngaros in the assemblies to the overall assembly sizes, and ii) we quantified the number of nucleotides in the PacBio reads that we could assign to either of the three fractions. The latter approach is less prone to assembly biases, such as overestimation of contributions for elements only present in one allele, or underestimation of contribution due to collapsed repeated copies, or elements not assembled because of low coverage. We aligned the PacBio reads to the assemblies using minimap2 v2.16 (-x map-pb) (46) and computed the coverage of the different genomic regions with samtools v1.9(47).

### Codon usage analysis

To analyze possible correlations between EMALE GC-content and the codon composition of their genes, we counted codons of all genes of complete EMALEs with a custom Perl script and visualized their distribution relative to their GC-content with a custom R script.

### Assignment of EMALE and Ngaro types

We first performed pairwise whole-genome comparisons of all 33 EMALEs plus the reference mavirus genome. Next, we concatenated the EMALE genomes according to their nearest sequence neighbors. We then plotted the resulting concatemer against itself with Gepard (48) using a word length of 10, and analyzed the similarity patterns. The dot plot-based classification was confirmed by phylogenetic analysis of virophage core genes. A similar approach was used for type assignment of Ngaro retrotransposons.

For partial EMALEs, we assigned types in an automated manner based on the highest cumulative blastx bitscores to typespecies EMALE genomes. EMALEs with cumulative bitscores below 100 were classified as “inconclusive”. For validation, the results were visualized with a beta version of gggenomes (https://github.com/thackl/gggenomes).

### Phylogenetic analysis of EMALE and Ngaro proteins

For EMALE core genes, multiple amino acid sequence alignments were constructed with MAFFT using the E-INS-i iterative refinement method (49). Alignments were manually inspected and trimmed to eliminate long insertions and regions of low sequence conservation. The best model and parameters for a maximum-likelihood phylogenetic reconstruction were estimated with modeltest-NG v0.1.6 (50) and the tree was computed with IQ-TREE v2.0 (ATPase: LG+I+G4, MCP: LG+R4+F, Penton: LG+I+G4+F, Protease: LG+R4+F, all: -B 1000) (51). The trees were visualized with ggtree v1.14.6 (52). For comparison, we also performed Bayesian inference analysis with MrBayes v3.1.2 using the following settings: rates=gamma, aamodelpr=mixed, number of generations=1 million (53).

For EMALE tyrosine recombinases, we generated a multiple amino acid sequence alignment with MAFFT v7.310 (--genafpair) (49) and an HMM profile (54) from the four sequences present in the type species genomes (EMALE05-EMALE08). We then identified closest relatives with jackhmmer on HmmerWeb v2.40.0 for 2 iterations (-E 1 --domE 1 --incE 0.001 --incdomE 0.001 --seqdb uniprotrefprot) (55), realigned the EMALE sequences with the 30 best hits (mafft --genafpair), and trimmed the alignments with trimAl (-automated1) (56). A maximum-likelihood tree was computed with FastTree v2.1.10 (57). The tree was visualized with ggtree v1.14.6 (52).

Phylogenies for the Ngaro YRs were generated the same way as for the EMALE YRs, however, only one iteration of jackhmmer was run, and only the top 20 database hits were included in the final tree.

### Comparative analysis of integration sites and their genomic context

For each of the 33 fully resolved EMALEs, we copied up to 10 kb (less than 10 kb if the EMALE was located within 10 kb of a contig border) of the 5’ and 3’ host sequences immediately flanking the TIR of the EMALE, and conducted blastn analyses on each of the four *C. burkhardae* genome assemblies with each flanking region separately. In case there was another EMALE located in the 10 kb flanking regions, the second EMALE was omitted from the BLAST search and only host sequence was included. After locating orthologous, and sometimes paralogous, sites in each host strain, we identified target site duplications by comparing empty and EMALE-containing alleles using pairwise sequence alignments of homologous sites. To analyze the genomic context of EMALEs for repetitive DNA, flanking regions were analyzed by dot plot analysis.

### Prediction of putative EMALE promoter motifs

We predicted putative promoter motifs in EMALE genomes by running MEME-suite v5.1.1 on all 100 bp upstream regions of all coding sequences. We identified the three highest-scoring motifs for each EMALE type individually. Putative motifs were further validated by analyzing their occurrences across all EMALE whole genomes, and motifs not consistently present in multiple intergenic regions were excluded.

### Validation of EMALE assemblies by PCR and Sanger sequencing

We designed primer pairs for selected EMALEs to generate overlapping, 700-1100 bp long PCR products. Primer sequences are available upon request. PCR products were obtained using 2 ng of genomic DNA template from *C. burkhardae* strain BVI, Cflag, E4-10, or RCC970 in 50 μl total volume containing 5 μl 10x Q5 Reaction Buffer (NEB, Frankfurt am Main, Germany), 0.5 U of Q5 High-Fidelity DNA Polymerase (NEB), 0.2 mM dNTPs, and 0.5 μM of each primer. Cycling conditions on a TGradient thermocycler (Biometra, Jena, Germany) consisted of: 30 s initial denaturation at 98 °C; 35 cycles of 10 s denaturation at 98 °C, 30 s annealing at 56-60 °C (depending on the melting temperature of the respective primers), and 45 s to 1 min extension at 72 °C; followed by a final extension time of 2 min at 72 °C. To check for correct product length and purity, 5 μl of each reaction were mixed with loading dye and analyzed on a 1% (w/v) agarose gel containing GelRed (VWR, Darmstadt, Germany). The remaining PCR mix was purified using a QIAquick PCR Purification Kit (Qiagen, Hilden, Germany) according to the manufacturer’s instructions. Sanger sequencing of PCR products was performed at Eurofins Scientific using the LightRun Tube service. Reads were trimmed, assembled, and mapped to their respective reference EMALEs in Sequencher™ software v5.2.2 (Gene Codes Corporation, Ann Arbor, MI, USA). Due to the presence of repetitive regions within and across EMALEs in a single host strain and resulting low-quality reads or failed PCRs, the Sanger assemblies typically did not cover the entire length of an EMALE.

### Effect of retrotransposon integration into EMALEs

To test if the insertion of a YR-retrotransposon triggered the degeneration of the targeted EMALE, we compared the lengths of conserved genes in Ngaro-containing EMALEs to those without transposon across all 138 EMALEs. Proteins length were obtained from the sequence files and plotted with a custom R script.

## Data availability

DNA sequences and genome annotations of endogenous virophages and Ngaro retrotransposons, as well as multiple sequence alignments used for phylogenetic reconstruction are available via https://github.com/thackl/cb-emales. Additional data items are deposited under https://doi.org/10.5281/zenodo.4632783

## Author Contributions

M.G.F. and T.H. conceived the project. K.B. and A.W. performed experiments. T.H. and M.G.F. analyzed, validated, and interpreted data. S.D. contributed to data analysis. T.H., M.G.F. and S.D. visualized data. T.H. developed software and performed computational analyses. M.G.F. supervised the study and wrote the original draft. M.G.F., T.H., and S.D. reviewed and edited the manuscript.

The authors declare no competing interests.

## Acknowledgments

This work was funded by the Max Planck Society and the Marine Microbiology Initiative of the Gordon and Betty Moore Foundation (Grant #5734). We thank I. Schlichting and C. Roome for support, and A. Koslová, J. Reinstein, C. Bellas and B. Müller for discussions.

