## Supplemental Figures for "Virophages and retrotransposons colonize the genomes of a heterotrophic flagellate"

**The supplementary information includes:**

Figures S1 to S13
Legends for Tables S1 to S3
Supplemental References

**Supplementary tables are provided as a supporting Excel file:**

Tables S1 to S3 (.xlsx)

### Supplementary Figures

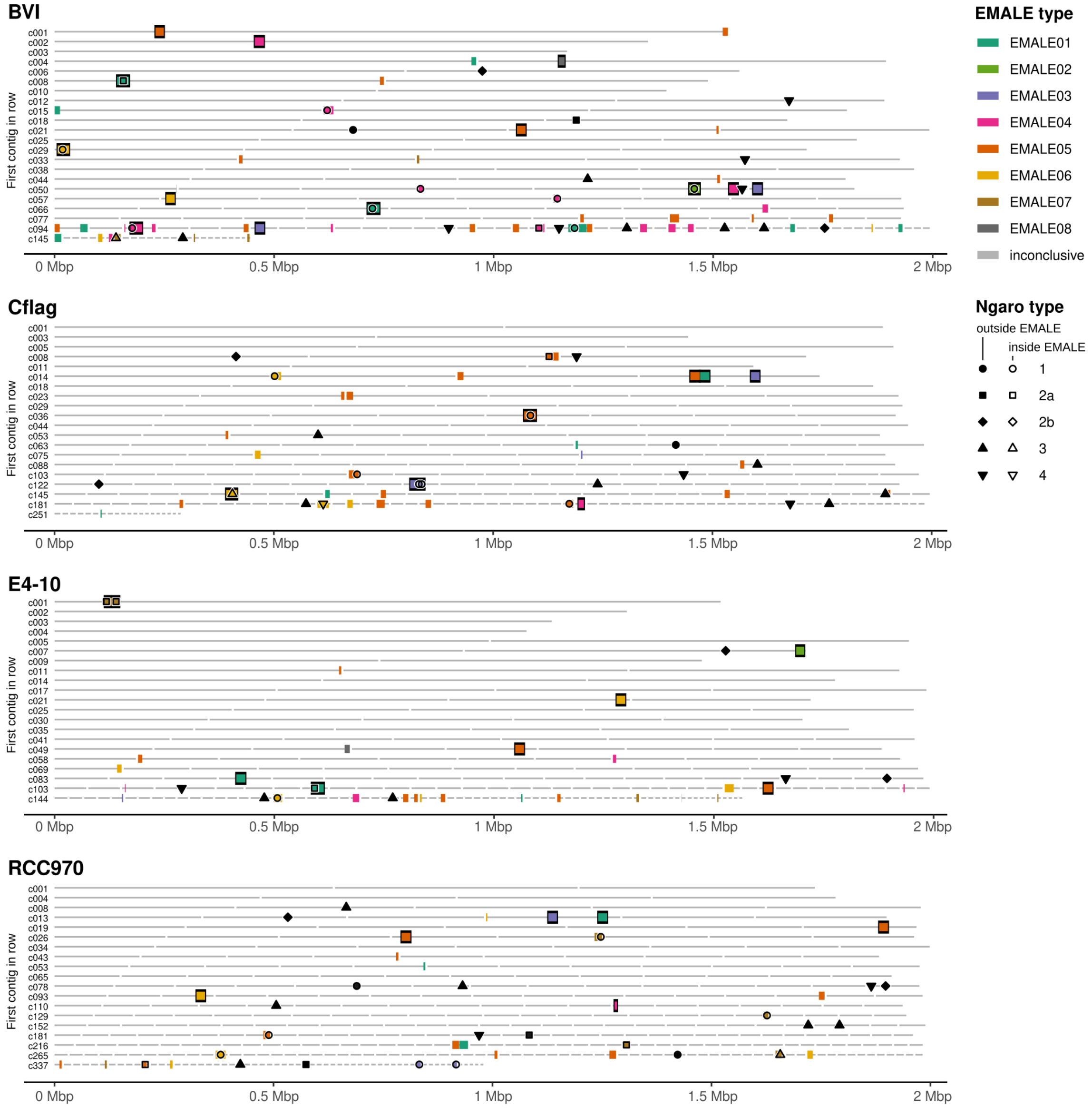


**Fig. S1. Distribution of EMALE and Ngaro retrotransposon integration sites in the four *Cafeteria burkhardae* genome assemblies.**
Location of partial and complete virophage genomes and Ngaro retrotransposons in the genome assemblies of four *C. burkhardae* strains. Horizontal lines represent contigs in order of decreasing length; colored boxes indicate the insertion sites of endogenous mavirus-like elements (EMALEs). Complete elements are framed in black. Ngaro retrotransposon insertion sites are marked by shapes corresponding to the different Ngaro types, open symbols indicate insertions in EMALEs.

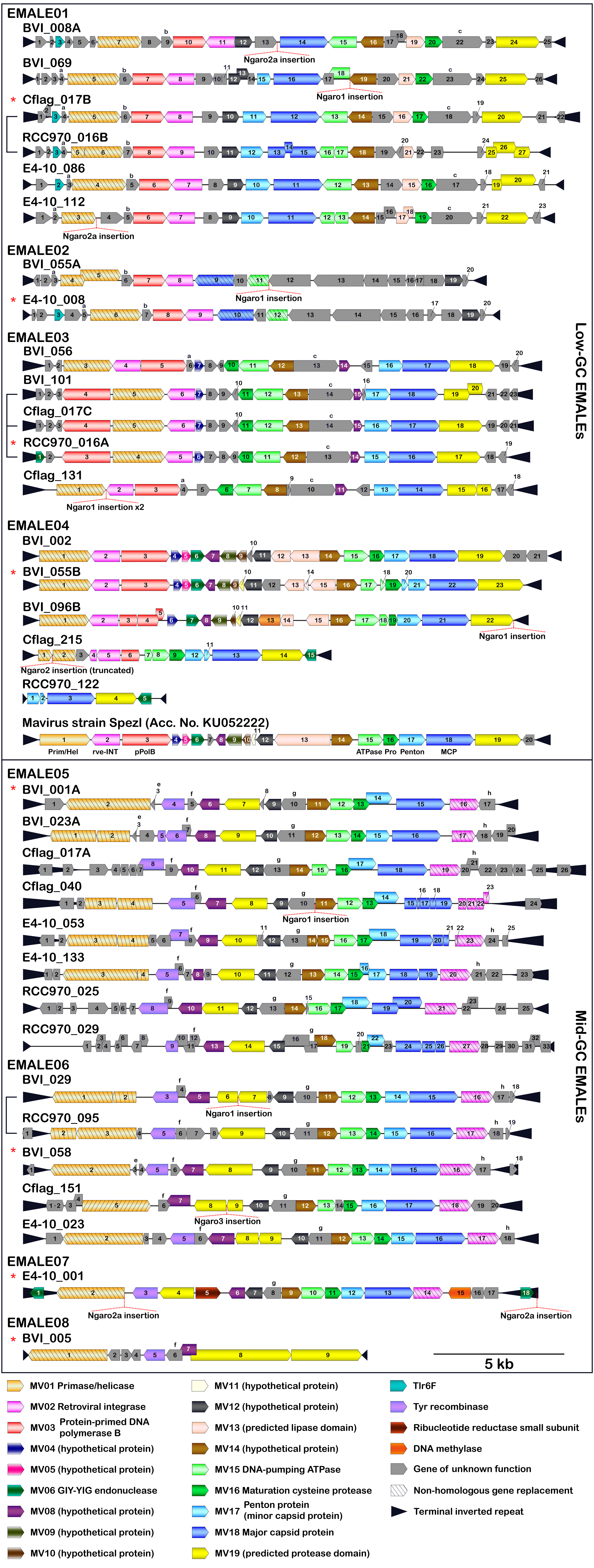
EMALE01
BVI_008A
Ngaro2a insertion
BVI_069
Ngaro1 insertion
Cflag_017B
RCC970_016B
E4-10_086
E4-10_112
Ngaro2a insertion
EMALE02
BVI_055A
Ngaro1 insertion
E4-10_008
EMALE03
BVI_056
BVI_101
Cflag_017C
RCC970_016A
Cflag_131
Ngaro1 insertion x2
EMALE04
BVI_002
BVI_055B
BVI_096B
Ngaro1 insertion
Cflag_215
Ngaro2 insertion (truncated)
RCC970_122
Mavirus strain Spezl (Acc. No. KU052222)
Prim/Hel
rve-INT
pPolB
ATPase
Pro
Penton
MCP
Low-GC EMALEs
EMALE05
BVI_001A
BVI_023A
Cflag_017A
Cflag_040
Ngaro1 insertion
E4-10_053
E4-10_133
RCC970_025
RCC970_029
EMALE06
BVI_029
Ngaro1 insertion
RCC970_095
BVI_058
Cflag_151
Ngaro3 insertion
E4-10_023
EMALE07
E4-10_001
Ngaro2a insertion
Ngaro2a insertion
EMALE08
BVI_005
5 kb
Mid-GC EMALEs
MV01 Primase/helicase
MV02 Retroviral integrase
MV03 Protein-primed DNA polymerase B
MV04 (hypothetical protein)
MV05 (hypothetical protein)
MV06 GIY-YIG endonuclease
MV08 (hypothetical protein)
MV09 (hypothetical protein)
MV10 (hypothetical protein)
MV11 (hypothetical protein)
MV12 (hypothetical protein)
MV13 (predicted lipase domain)
MV14 (hypothetical protein)
MV15 DNA-pumping ATPase
MV16 Maturation cysteine protease
MV17 Penton protein (minor capsid protein)
MV18 Major capsid protein
MV19 (predicted protease domain)
Tlr6F
Tyr recombinase
Ribucleotide reductase small subunit
DNA methylase
Gene of unknown function
Non-homologous gene replacement
Terminal inverted repeat

**Fig. S2 (previous page). Coding capacity of 33 completely assembled EMALEs in *C. burkhardae*.**

Shown are genome diagrams for 33 EMALEs in four *C. burkhardae* strains (BVI, Cflag, E4-10, RCC970).The reference mavirus genome with genes MV01-MV20 is included for comparison. EMALE identifiers consist of host strain name, followed by contig number and sometimes letters to distinguish between several EMALEs on the same contig. Directional boxes indicate ORFs in the respective orientation. Homologous genes that are present in mavirus or have a predicted function are shown in color. Other homologous ORFs are denoted by lowercase letters. Black triangles represent terminal inverted repeats (TIRs). Brackets indicate EMALEs with homologous integration sites in different host strains. Ngaro retrotransposon insertions are shown when present. Asterisks denote type elements as shown in Fig. 3.

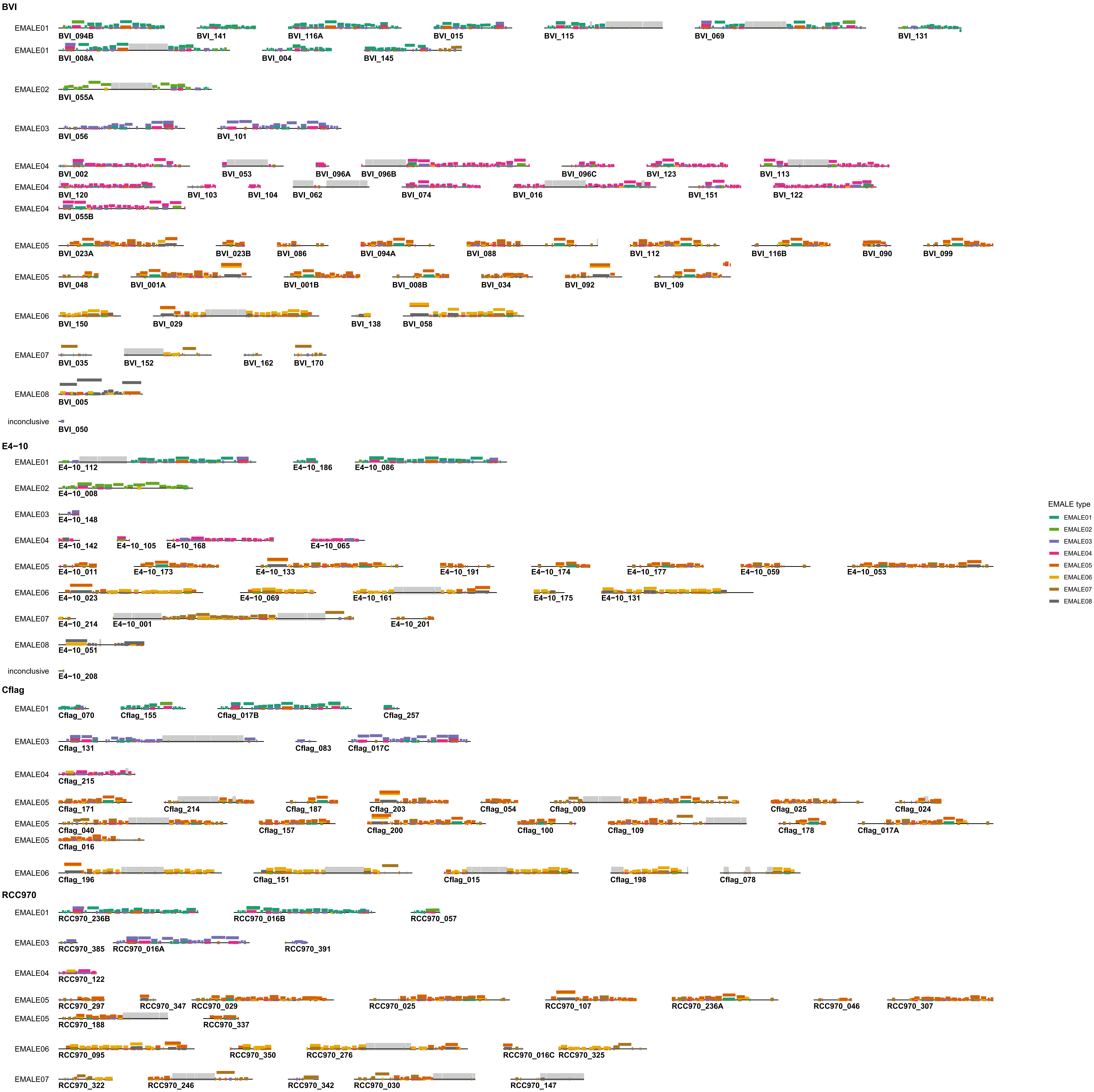
BVI
EMALE01
BVI_094B
BVI_141
BVI_116A
BVI_015
BVI_115
BVI_069
BVI_131
EMALE01
BVI_008A
BVI_004
BVI_145
EMALE02
BVI_055A
EMALE03
BVI_056
BVI_101
EMALE04
BVI_002
BVI_053
BVI_096A
BVI_096B
BVI_096C
BVI_123
BVI_113
EMALE04
BVI_120
BVI_103
BVI_104
BVI_062
BVI_074
BVI_016
BVI_151
BVI_122
EMALE04
BVI_055B
EMALE05
BVI_023A
BVI_023B
BVI_086
BVI_094A
BVI_088
BVI_112
BVI_116B
BVI_090
BVI_099
EMALE05
BVI_048
BVI_001A
BVI_001B
BVI_008B
BVI_034
BVI_092
BVI_109
EMALE06
BVI_150
BVI_029
BVI_138
BVI_058
EMALE07
BVI_035
BVI_152
BVI_162
BVI_170
EMALE08
BVI_005
inconclusive
BVI_050
E4−10
EMALE01
E4−10_112
E4−10_186
E4−10_086
EMALE02
E4−10_008
EMALE03
E4−10_148
EMALE04
E4−10_142
E4−10_105
E4−10_168
E4−10_065
EMALE05
E4−10_011
E4−10_173
E4−10_133
E4−10_191
E4−10_174
E4−10_177
E4−10_059
E4−10_053
EMALE06
E4−10_023
E4−10_069
E4−10_161
E4−10_175
E4−10_131
EMALE07
E4−10_214
E4−10_001
E4−10_201
EMALE08
E4−10_051
inconclusive
E4−10_208
Cflag
EMALE01
Cflag_070
Cflag_155
Cflag_017B
Cflag_257
EMALE03
Cflag_131
Cflag_083
Cflag_017C
EMALE04
Cflag_215
EMALE05
Cflag_171
Cflag_214
Cflag_187
Cflag_203
Cflag_054
Cflag_009
Cflag_025
Cflag_024
EMALE05
Cflag_040
Cflag_157
Cflag_200
Cflag_100
Cflag_109
Cflag_178
Cflag_017A
EMALE05
Cflag_016
EMALE06
Cflag_196
Cflag_151
Cflag_015
Cflag_198
Cflag_078
RCC970
EMALE01
RCC970_236B
RCC970_016B
RCC970_057
EMALE03
RCC970_385
RCC970_016A
RCC970_391
EMALE04
RCC970_122
EMALE05
RCC970_297
RCC970_347
RCC970_029
RCC970_025
RCC970_107
RCC970_236A
RCC970_046
RCC970_307
EMALE05
RCC970_188
RCC970_337
EMALE06
RCC970_095
RCC970_350
RCC970_276
RCC970_016C
RCC970_325
EMALE07
RCC970_322
RCC970_246
RCC970_342
RCC970_030
RCC970_147
EMALE type
EMALE01
EMALE02
EMALE03
EMALE04
EMALE05
EMALE06
EMALE07
EMALE08

**Fig. S3 (previous page). Type-assignment for incomplete EMALEs.**
For partial EMALEs, we assigned types in an automated manner based on blastx hits to type species EMALEs. For each EMALE, blast hits to type species proteins along the genome are shown. Hit positions on the y-axis reflect bitscores, with the highest-scoring hits plotted at the top. Integrated Ngaro retrotransposons are shown as grey boxes.

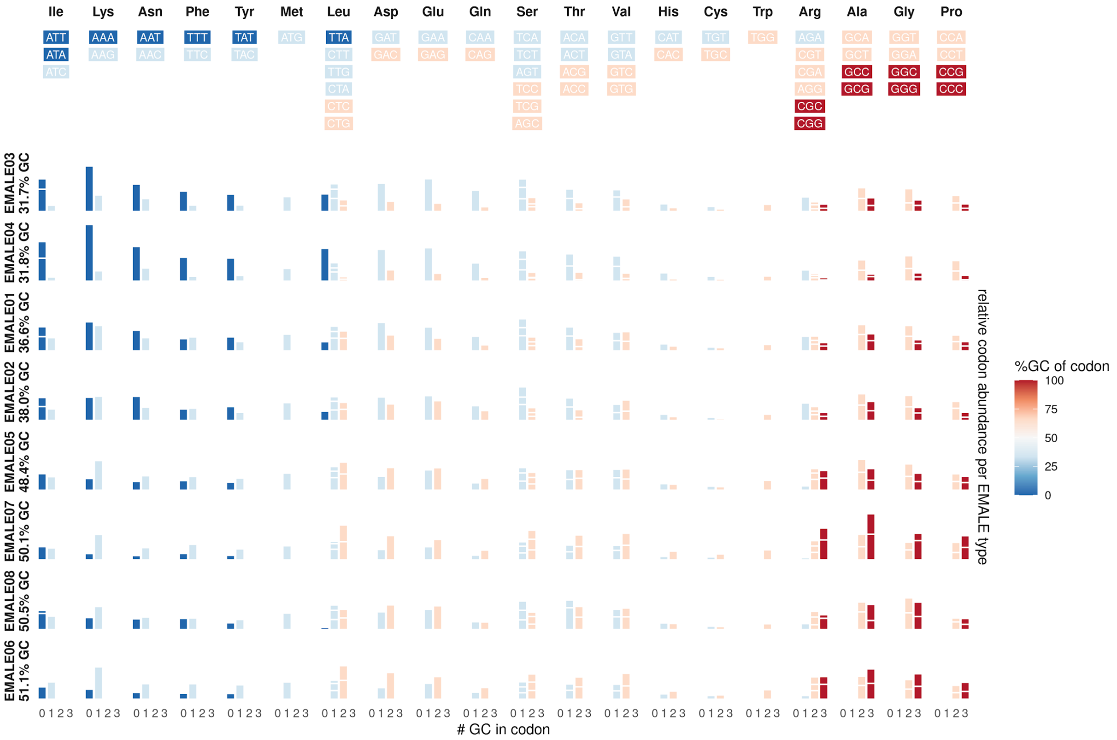


**Fig. S4. Codon usage with respect to GC-content in different EMALE types.**
Each row represents one of the eight EMALE types, ordered by decreasing GC-content. Each column represents a single amino acid. The different codons coding for each amino acid are indicated above the plot, color-coded according to their GC-content. The histograms indicate the relative contributions of each codon with a certain GC-content to the overall amino acid composition of each EMALE. Codons with identical GC-content encoding the same amino acid are represented by stacked bars separated by white lines.

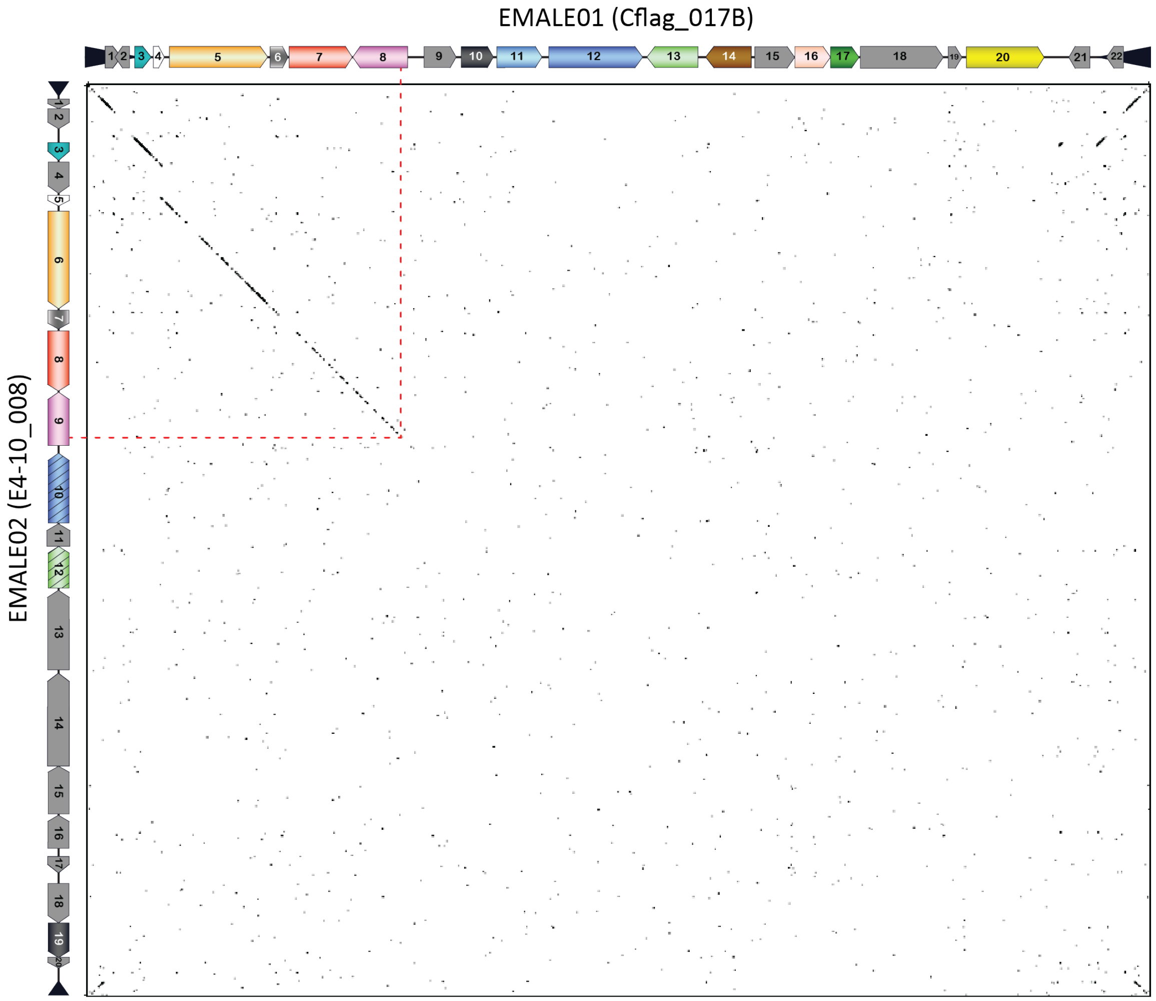


**Fig. S5. Partial synteny between EMALE01 and EMALE02.**
DNA dot plot analysis of EMALE01 Cflag_017B and EMALE02 E4-10_008 showing predicted genes along the axes. The synteny ends within the rve-INT gene, which represents the presumed recombination site (red dotted line). For the ORF color legend, see Figs. 3, S2.

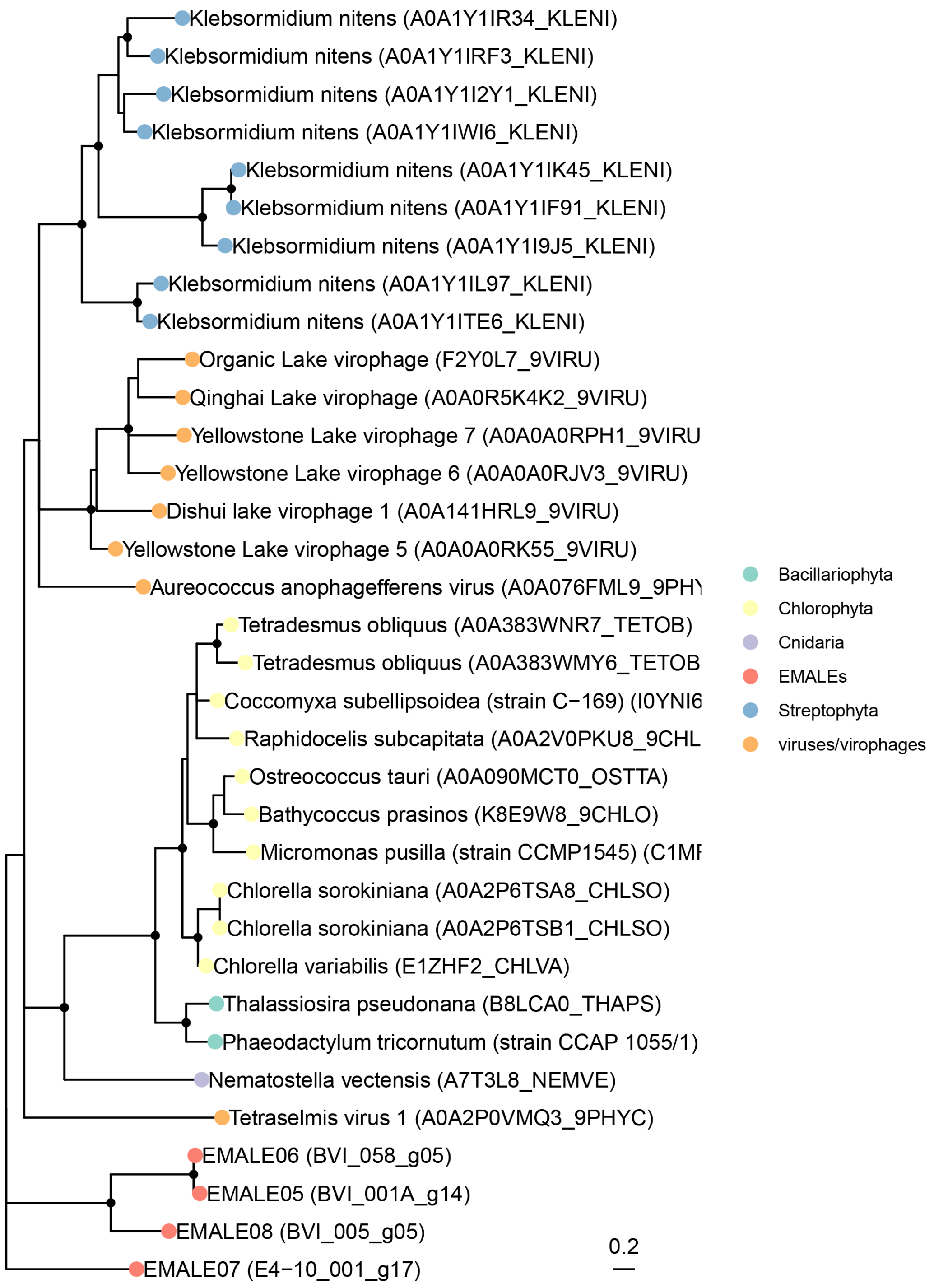


**Fig. S6. Maximum likelihood reconstruction of EMALE tyrosine recombinase phylogenies.** Shown is an unrooted ML phylogenetic tree of tyrosine recombinases comparing EMALE proteins (red dots) to their most similar sequences in UniProt (color-coded at the phylum level). Nodes with <50% bootstrap support were collapsed; nodes with >80% bootstrap support are marked by solid black dots.

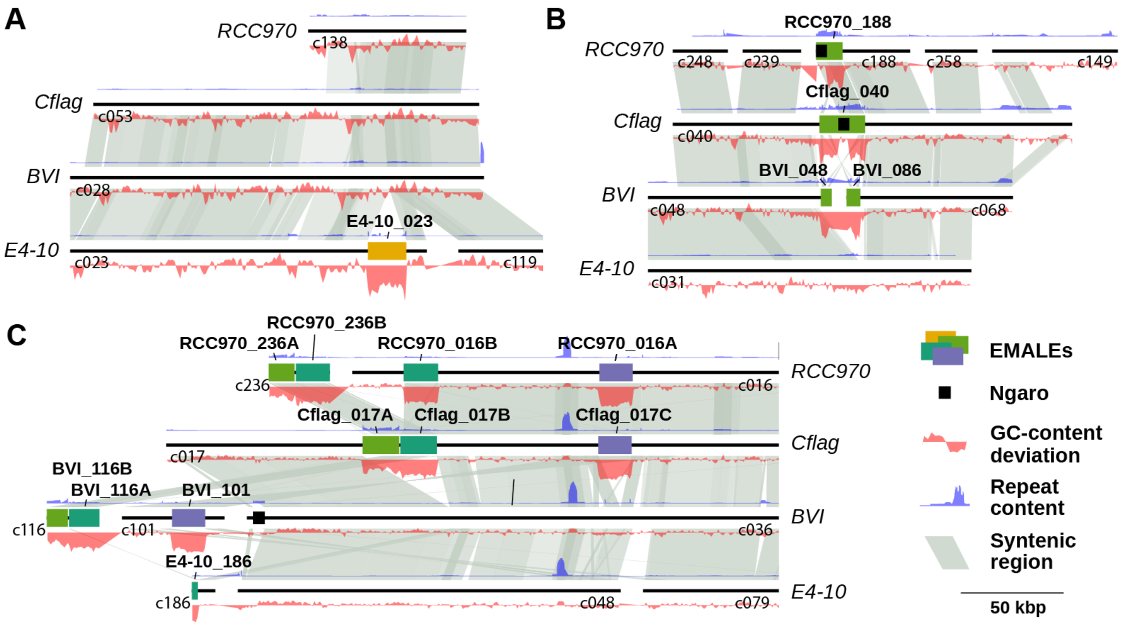


**Fig. S7. Unique and orthologous EMALE integration loci among four *Cafeteria* strains.** Synteny plots of three genomic loci illustrate different scenarios of EMALE conservation in *C. burkhardae*. Homologous DNA regions are connected by grey shadings. Peaks in the blue curve indicate repetitive regions, red curves represent GC-content. (A) EMALE E4-10_023 represents a unique integration site in host strain E4-10. Syntenic regions in other host strains are well resolved and devoid of EMALEs. (B) Homologs of EMALE Cflag_040 are found in orthologous loci in host strains RCC970 and BVI. The Ngaro retrotransposon in this EMALE apparently caused assembly problems, resulting in premature termination of RCC970 contig 188 and splitting the EMALE onto two contigs in BVI. (C) Comparative analysis of the three EMALEs on Cflag contig 17 reveals a complex situation. The double EMALE Cflag_017A/B has homologs in BVI and RCC970, albeit as partial elements on short contigs. Most of RCC970 contig 16 is syntenic to Cflag contig 17, except for the shorter flanking region of EMALE RCC970_016B, which is likely caused by mis-assembly of the RCC970 contig at the double-EMALE transition. In the BVI assembly, the flanking regions of EMALE Cflag_017C are present twice, on the EMALE-containing contig 101 and on the EMALE-free contig 36, suggesting a heterozygous condition in BVI.

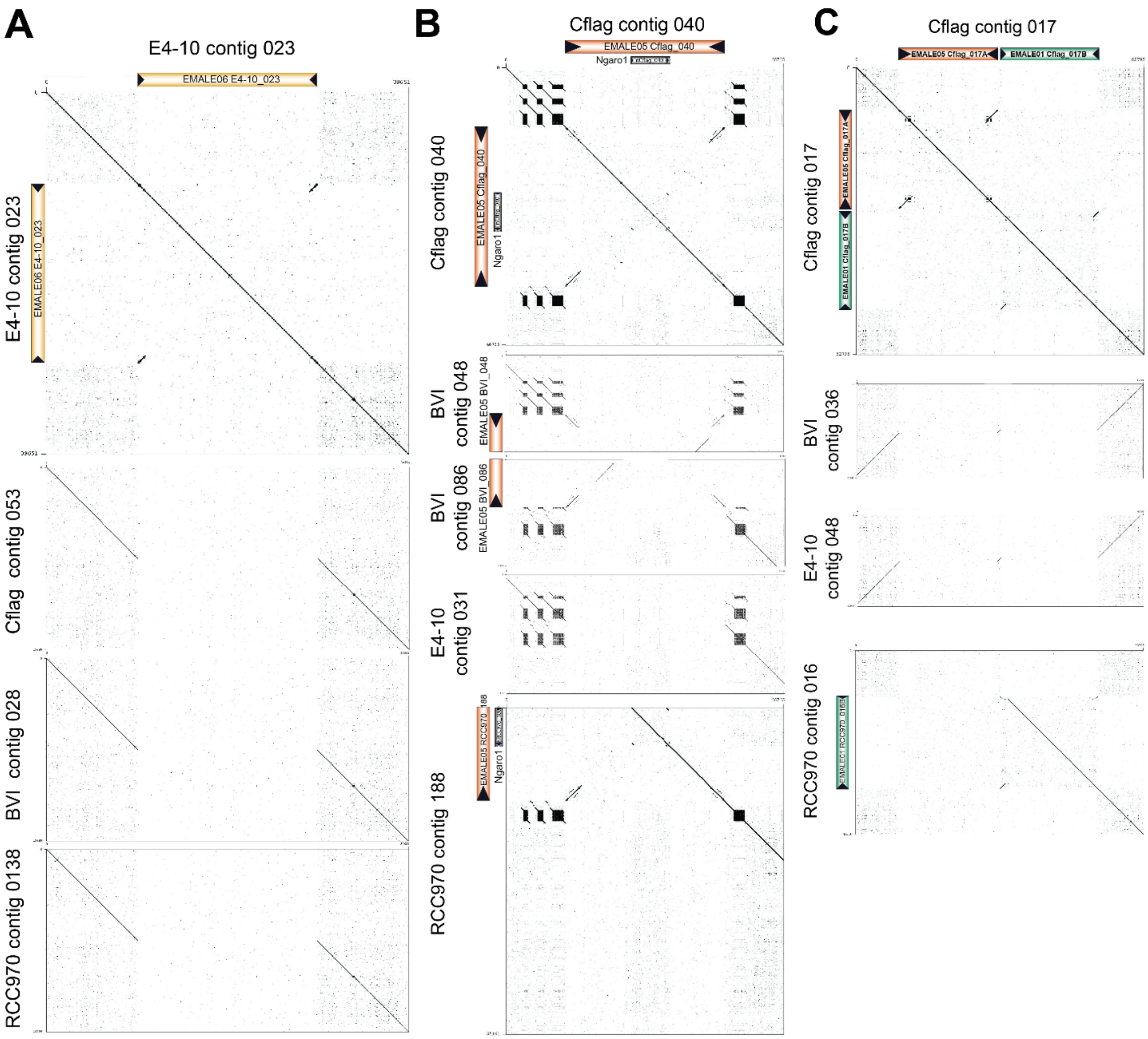


**Fig. S8. DNA dot plots of selected EMALE loci as shown in Fig. S7.**

Shown are DNA dot plots of EMALE(s) including 10 kb of flanking host DNA versus itself and versus syntenic regions in other host strains. Black triangles represent EMALE TIRs. (A) EMALE E4-10_023 is integrated in non- repetitive host DNA and represents a unique insertion in strain E4-10, with EMALE-free loci in the other three strains. (B) EMALE Cflag_040 is integrated in a cluster of complex host repeats and has homologous integration sites in strains BVI and RCC970. (C) The double EMALE Cflag_017A/B likely caused mis-assembly in strain RCC970.

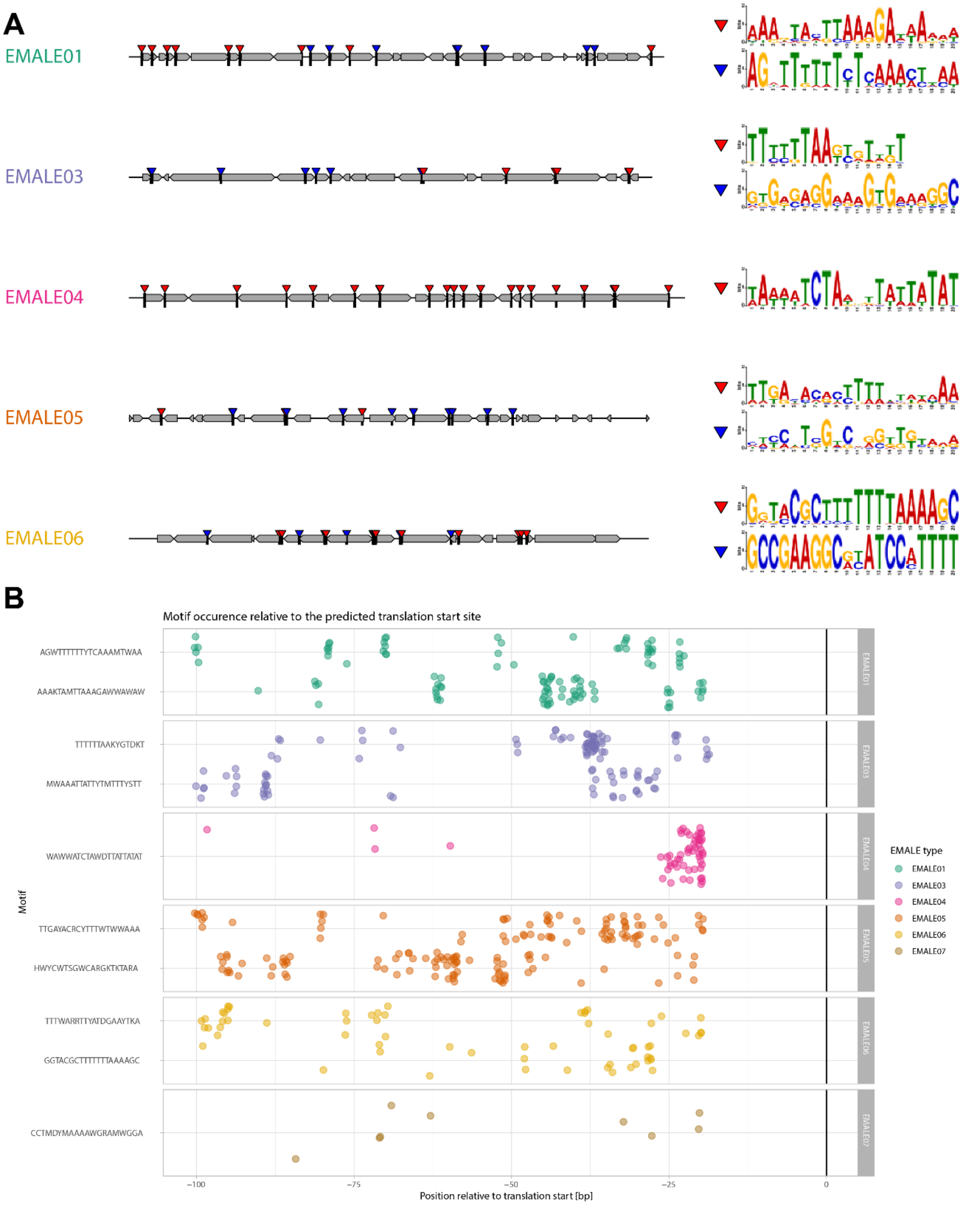


**Fig. S9. Putative promoter motifs in EMALE genomes.**
(A) Sequence logos of high-scoring motifs predicted with MEME in immediate upstream regions of coding sequences and their positions in EMALE genomes. Character height at each position of a sequence logo corresponds to the frequency of the respective nucleotide at that position. (B) EMALE promoter motif occurrences relative to predicted translation start sites. Each dot corresponds to the start of a predicted promoter motif plotted relative to the ATG start codon of the downstream gene. Motifs are grouped by the EMALE type in which they were initially predicted.

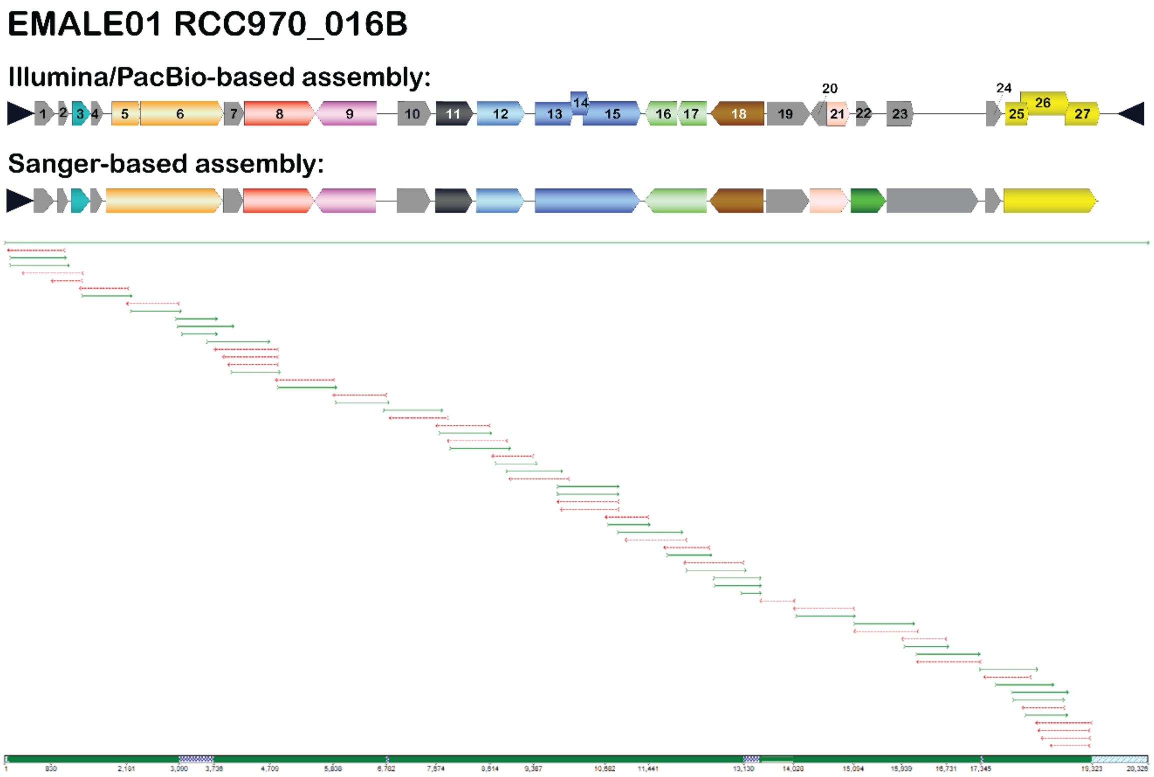


**Fig. S10. Correction of Illumina/PacBio-based assemblies by PCR and Sanger sequencing.** EMALE01 RCC970_016B was re-evaluated by PCR analysis and subsequent Sanger sequencing of PCR products. The resulting assembly is compared to the Illumina/PacBio-based assembly (top). The bottom part of the figure shows a Sequencher™ screenshot of the assembled Sanger reads. The long green arrow represents the Illumina/PacBio sequence, the shorter green and red arrows represent individual Sanger reads.

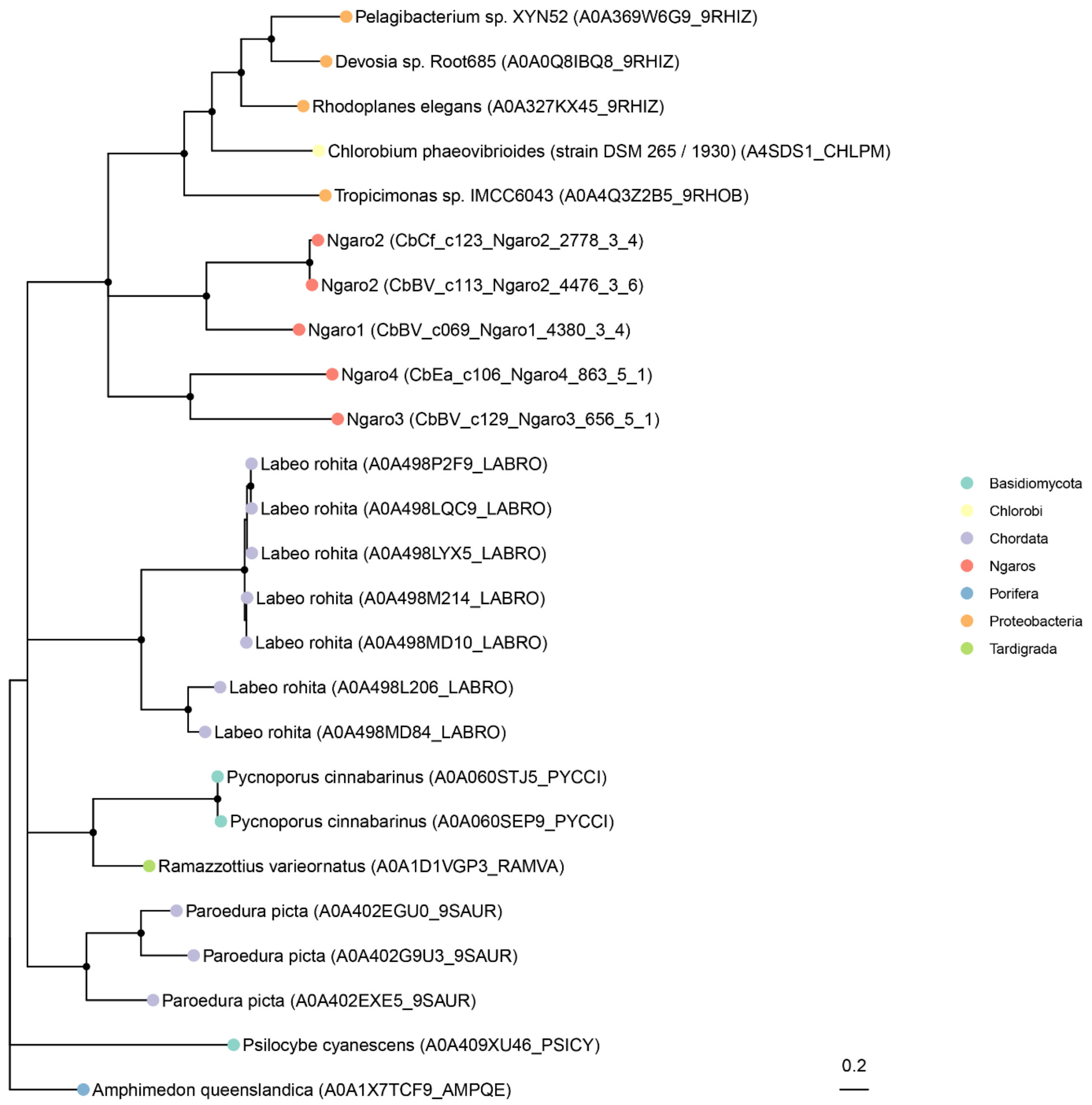


**Fig. S11. Phylogenetic placement of Ngaro tyrosine recombinases.**
Shown is an unrooted maximum likelihood phylogenetic tree of tyrosine recombinases comparing *Cafeteria* Ngaro-encoded proteins (red dots) with their most similar sequences in UniProt (color-coded at the phylum level). Nodes with <50% bootstrap support were collapsed; nodes with >80% bootstrap support are marked by black dots. The distribution across a very broad range of phyla (fish, fungi, sponges, tardigrades, and bacteria) indicates that this integrase is likely associated with highly mobile transposons that have broad host ranges, confirming previous studies on these elements (1).

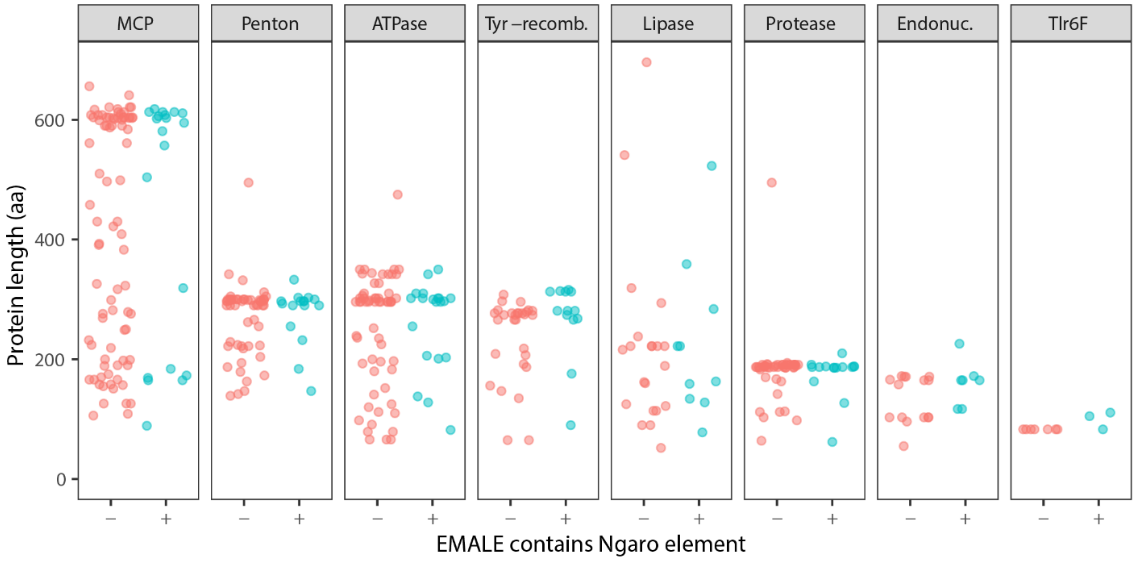


**Fig. S12. Protein length distributions in EMALEs with and without retrotransposons.** Integration of Ngaro retrotransposons might lead to the inactivation of EMALEs, thereby promoting their degeneration and the pseudogenization of their genes. To test this hypothesis, we compared the predicted protein lengths for conserved genes in EMALEs without (red) and with transposons (blue). Gene length here serves as a proxy for pseudogenization because this process leads to the emergence of premature stop codons and frameshifts, which in turn leads to shorter annotated genes.

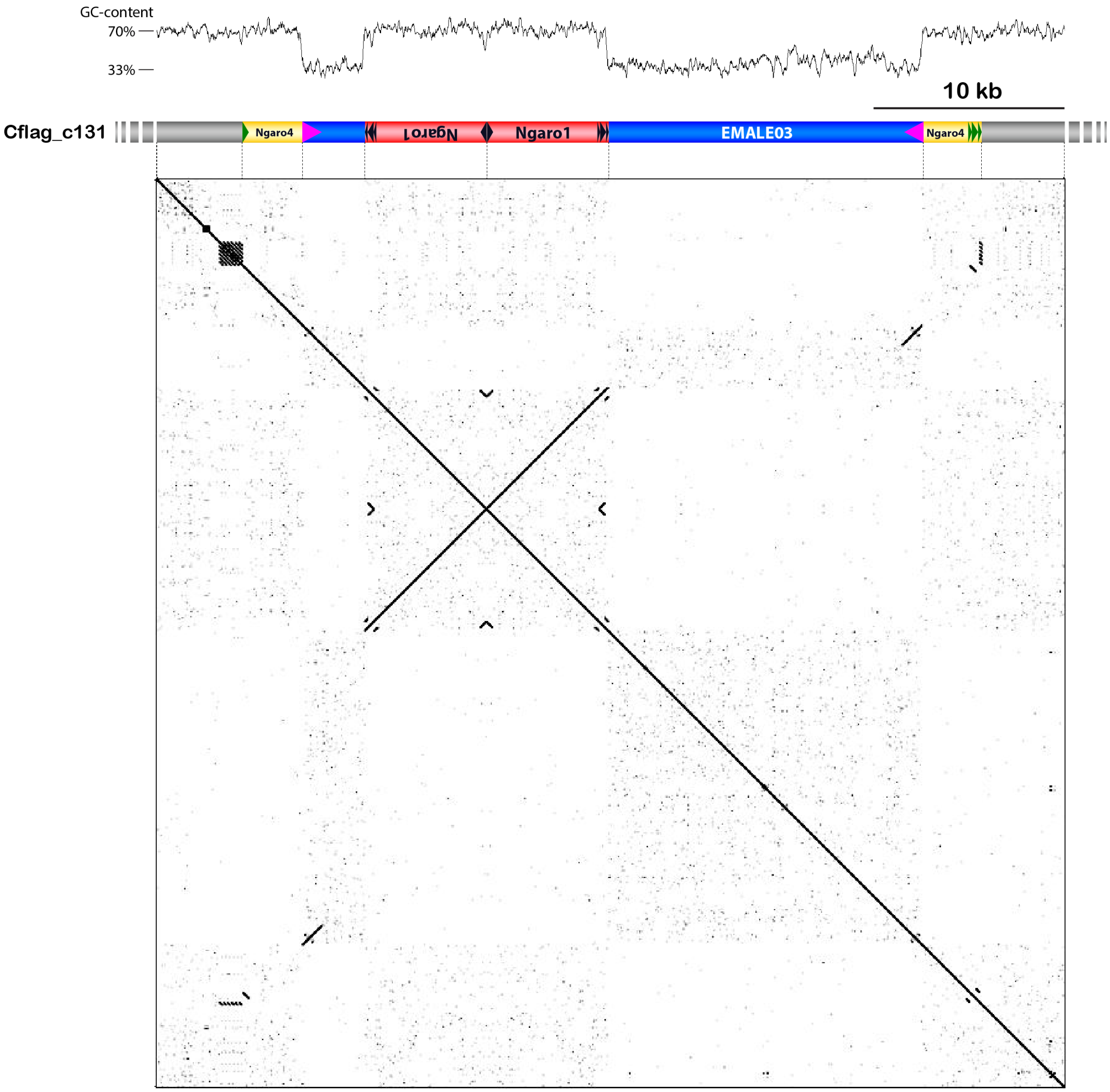


**Fig. S13. Nested integration scenario involving one EMALE and three Ngaro retrotransposons.**

EMALE03 Cflag_131 (blue) is inserted into a type 4 Ngaro retrotransposon (yellow) and contains two insertions of type 1 Ngaros (red). The nested integration scenario is depicted by a GC-content graph (top), a graphic representation of the respective genomic region on contig 131 of *C. burkhardae* strain Cflag (middle), and a self-versus-self DNA dot plot (bottom). Host sequence is shown in grey; terminal repeats are represented by colored triangles.

**Table S1 (supporting .xlsx file). EMALE statistics.** This dataset contains information on each of the 138 EMALEs in *C. burkhardae*, including their exact location in the host assembly, length, presence of terminal inverted repeats, type score, and Ngaro insertions.

**Table S2 (supporting .xlsx file). EMALE integration sites.** This dataset lists information for each of the 33 fully assembled EMALEs regarding orthologous integration loci in all four host strains, target site duplications, and host genomic context of the integration loci.

**Table S3 (supporting .xlsx file). Ngaro statistics.** This dataset contains information for 80 Ngaro retrotransposons identified in the four *C. burkhardae* assemblies, including their exact location in the host assembly, length, type, and insertion locus (EMALE or eukaryotic chromatin).
